# Somatic aging pathways regulate reproductive plasticity in *Caenorhabditis elegans*

**DOI:** 10.1101/673764

**Authors:** Maria C. Ow, Alexandra M. Nichitean, Sarah E. Hall

## Abstract

Early life stress of an animal often results in changes in gene expression that correspond with changes in their adult phenotype. In the nematode *C. elegans*, starvation during early larval stages promotes entry into a non-feeding, stress-resistant stage named dauer until environmental conditions improve. Here we show that the endocrine signaling attributed to the somatic aging pathways in *C. elegans* adults lacking a functional germline also regulate the reproductive phenotypes of adults that have experienced dauer as a result of early life starvation. Postdauer adults modulate their fatty acid metabolism in order to re-allocate fat reserves in a manner benefitting their progeny at the expense of the parental somatic fat reservoir. Our results also show that the metabolic plasticity in postdauer animals and the inheritance of ancestral starvation memory in the progeny are a result of crosstalk between somatic and reproductive tissues mediated by the HRDE-1 nuclear RNAi Argonaute.

## Introduction

Evidence indicating that experiences during early development affect behavior and physiology in a stress-specific manner later in life is abundant throughout the animal kingdom (Telang and Wells, 2004; Weaver et al. 2004; Binder et al. 2008; Pellegroms et al. 2009; van Abeelen et al. 2012; Zhao et al. 2014; Canario et al. 2017; Dantzer et al. 2019; Vitikainen et al. 2019). Epidemiological studies and studies using mammalian animal models have supported the “thrifty” phenotype hypothesis which proposes that fetal or postnatal malnutrition results in the increased risk for metabolic disorders in the offspring (Neel, 1962; Hales and Barker, 1992; Vaag et al. 2012; Smith and Ryckman, 2015). For instance, individuals exposed to the WWII Dutch Hunger Winter during gestation had lower glucose tolerance and increased risk of obesity, diabetes, and cardiovascular diseases in adulthood compared to siblings born before the famine. In addition, the increased propensity to develop metabolic disorders was inherited for two generations (Painter et al. 2008; Lumey et al. 2011; Veenendaal et al. 2013). Thus, understanding the relationship between fetal malnutrition and the increased susceptibility to metabolic disorders in adulthood is critical for development of intervention and therapeutic strategies.

The nematode *C. elegans* makes a critical decision regarding its developmental trajectory based on the environmental conditions experienced during its first larval stage (L1). If conditions are poor (e.g. low food availability, crowding, or high temperatures), decreased insulin and TGF-β signaling in L1 larvae promote entry into an alternative, stress-resistant, non-aging, diapause stage named dauer. Once conditions improve, dauer larvae resume development as postdauer L4 larvae and continue through reproductive adulthood as postdauer adults (Cassada and Russell, 1975). Alternatively, if conditions are favorable, L1 larvae proceed through additional larval stages (L2-L4) until reaching reproductive adulthood (control adults) (Sulston and Horvitz, 1977). Although postdauer adults are morphologically similar to control adults, we previously showed that postdauer adults retained a cellular memory of their early life experience that resulted in genome-wide changes in their chromatin, transcriptome, and life history traits (Hall et al. 2010, 2013; Ow et al. 2018). Remarkably, postdauer adults also encoded the nature of their L1 environmental experience and gauged their adult reproductive phenotypes and genome-wide gene expression based on this memory (Ow et al. 2018). Postdauer adults that experienced crowding or high pheromone conditions exhibited increased fertility and upregulated expression of genes involved in reproduction relative to control adults that never experienced crowding. In contrast, postdauer adults that experienced starvation (PD_Stv_) exhibited decreased fertility and an enrichment in somatic gene expression compared to control adults that never experienced starvation (CON_Stv_) (Ow et al. 2018). Moreover, the changes in fertility and somatic gene expression in PD_stv_ adults required a functional germ line (Ow et al. 2018).

The prerequisite of crosstalk between somatic and reproductive tissues for the postdauer reproductive phenotypes is also a key regulatory feature governing adult lifespan and stress response (Kenyon 2010, 2010b). In *C. elegans*, endocrine signaling has emerged as one of the principal pathways extending the lifespan of animals lacking a germ line due to either ablation of germ line precursor cells or a mutation in the *glp-1*/Notch receptor gene. The two main effectors of endocrine signaling, the FOXO transcription factor DAF-16 and the nuclear hormone receptor (NHR) DAF-12, are required for the increased lifespan of germline-less animals and are regulated by the physiological state of the animal (Hsin and Kenyon, 1999; Kenyon 2010, 2010b; Murphy and Hu, 2013). When an animal experiences reproductive stress, including sterility, DAF-16 is dephosphorylated and translocated to the nucleus where it can alter target gene expression to promote the extended longevity of germ line-less animals (Kenyon 2010, 2010b; Murphy and Hu, 2013). DAF-12, a homolog of the mammalian vitamin D receptor, binds to bile acid-like steroid ligands (dafachronic acids or DA) to promote the expression of genes involved in reproduction and growth in favorable conditions (Antebi 2014). In *glp-1* mutants, the Δ^7^ form of DA (Δ^7^-DA) is four-fold increased compared to wild-type and promotes DAF-16 nuclear localization (Berman and Kenyon, 2006; Shen et al. 2012). One of the consequences of the DAF-16 and DAF-12 dependent endocrine signaling in *glp-1* mutants is a significant increase in stored intestinal fat, which allows for somatic maintenance and prolonged longevity in the absence of germline development (Wang et al. 2008).

In this study, we show that steroid hormone signaling, reproductive longevity signaling, and nuclear hormone receptors contribute to the decreased fertility of postdauer animals that experienced early life starvation by regulating fatty acid metabolism. The reproductive plasticity of PD_stv_ adults is a result of crosstalk between somatic and reproductive tissues, the effect of which is an increase in lipid metabolic pathway function in an animal that has experienced dauer, resulting in decreased fertility but endowing metabolic robustness to their progeny. Thus, the pathways that bestow increased lipid storage and extended longevity in a germline-less animal function to promote reproduction in a postdauer animal that experienced early-life starvation. We also show that the signals between the soma and the germ line are dependent on HRDE-1, a germline-specific nuclear RNAi argonaute. Given the principal role of HRDE-1 in RNAi-mediated transgenerational inheritance, this pathway may transmit the ancestral starvation memory to an ensuing generation and provide the necessary hardiness to survive future famine.

## Results

### Dafachronic acid-dependent DAF-12 signaling is required for decreased fecundity after starvation-induced dauer formation

Given that endocrine signaling across tissues is a prominent feature of reproductive longevity, we examined whether wild-type PD_Stv_ adults shared any gene expression signatures with animals lacking a functional germ line. In *glp-1* mutants, increased longevity is dependent on TOR (target of rapamycin) signaling, DAF-16/FOXO gene regulation, steroid hormone signaling, and fatty acid metabolism regulation (Lapierre and Hansen, 2012). With the exception of TOR signaling, we found significant gene expression changes in PD_Stv_ adults of key genes in each of these regulatory pathways.

In the steroid signaling pathway, dafachronic acid (DA) biosynthesis requires the cytochrome P450 DAF-9, the Reiske-like oxygenase DAF-36, and the hydroxysteroid dehydrogenase HSD-1 (Fig. 1A) (Gerisch et al. 2004; Rottiers et al. 2006; Patel et al. 2008; Mahanti et al. 2014). In animals lacking a functional germ line, the levels of *daf-36* mRNA and the Δ^7^ form of DA (Δ^7^-DA) are significantly increased compared to wild-type (Shen et al. 2012). Interestingly, we found that in wild-type PD_Stv_ adults with a germ line, *daf-36* mRNA is also increased 3-fold (*p* = 3.25e-04; FDR = 0.01) compared to control adults (Ow et al. 2018). To investigate whether DA signaling plays a role in mediating reproductive plasticity as a result of early life experience, we asked whether mutations in DA biosynthesis genes would affect the reduced brood size observed in PD_Stv_ adults. We found that *daf-9(rh50)* and *daf-36(k114)* strains, but not *hsd-1(mg433)*, exhibited a significant increase in brood size in PD_Stv_ adults, opposite of what we observed in wild-type animals (Fig. 1B; Supplemental Table S1).

**Figure 1.**
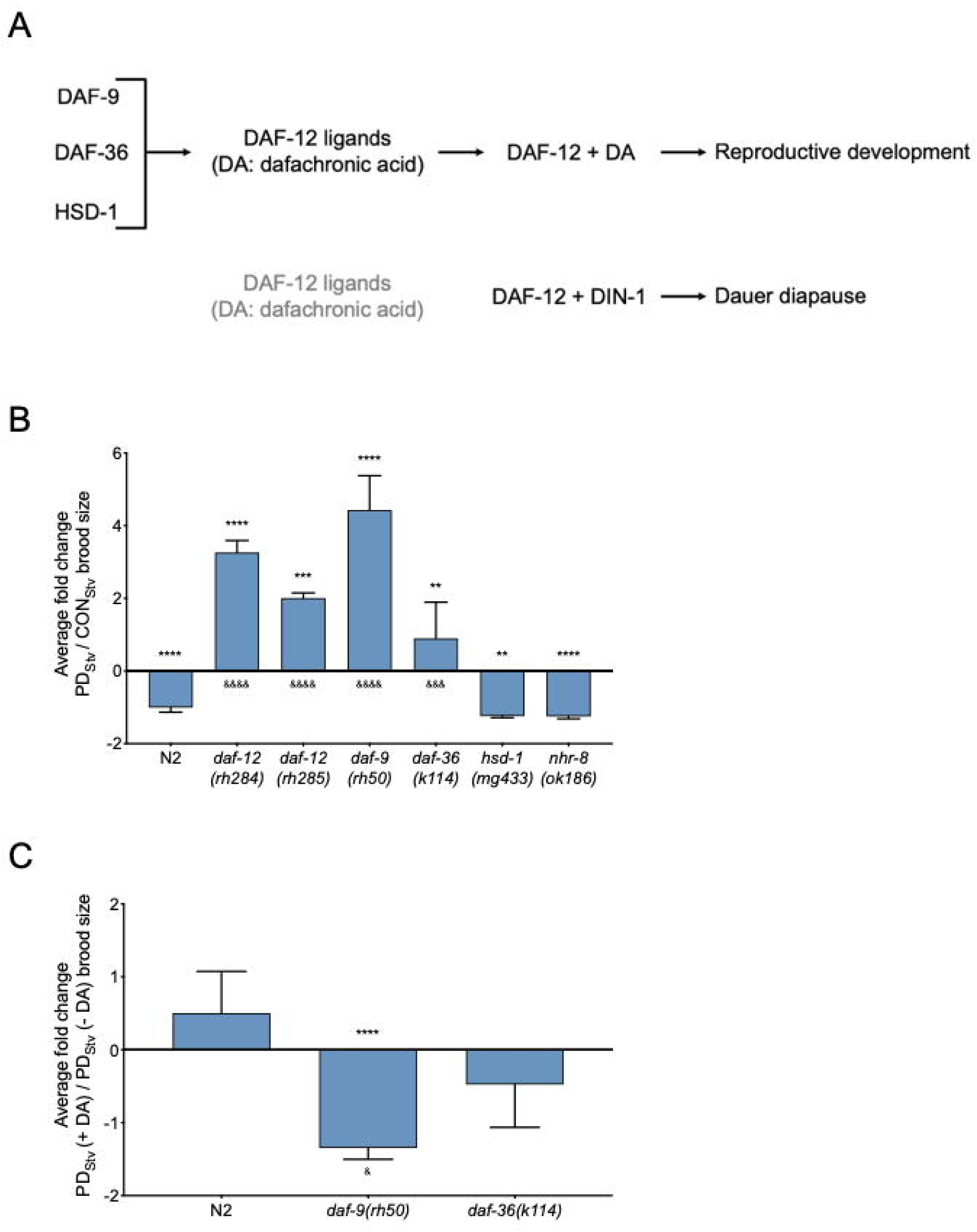
Adult reproductive plasticity is dependent on DAF-12 steroid signaling. (*A*) Model of DAF-12 NHR regulation of development. See text for details. (*B*) Brood size comparison of PD_Stv_ relative to CON_Stv_ in wild-type N2 and mutant strains represented as the mean ± S.E.M. ** *p* < 0.01, *** *p* < 0.001, **** *p* < 0.0001; Student’s *t*-test; comparison of PD_Stv_ to CON_Stv_ of a given genotype. ^&&&^ *p* < 0.001, ^&&&&^ *p* < 0.0001; one-way ANOVA with Fisher’s LSD test; comparison of PD_Stv_/CON_Stv_ brood size between mutant and wild-type N2. Additional brood size data is provided in Supplemental Table S1. (*C*) Brood size comparison of PD_Stv_ grown in the presence of 40 nM Δ^7^-DA relative to control, represented as the mean ± S.E.M. N= 4 trials, **** *p* < 0.0001 (Student’s *t*-test) indicates the comparison between PD_Stv_ with Δ^7^-DA and control. ^&^ *p* < 0.05 (one-way ANOVA with Fisher’s LSD *post hoc* test) refers to the comparison of brood size fold change between *daf-9* and wild-type N2. Additional brood size data can be found in Supplemental Table S2.

We next asked whether the steroid signaling that contributed to the reduced fertility of PD_Stv_ adults acted through related NHRs, DAF-12 and NHR-8. Since null mutants of *daf-12* are dauer defective, we used two *daf-12* alleles, *rh284* (Class 5 mutant) and *rh285* (Class 4 mutant), with lesions in the ligand binding domain (LBD) that affect steroid signaling activity but otherwise have nearly normal dauer phenotype (Antebi et al. 2000). Surprisingly, *daf-12(rh284)* and *daf-12(rh285)* exhibited a significant increase in brood size, similar to that observed for the DA biosynthesis mutants (Fig. 1B; Supplemental Table S1). These results suggest that DA-dependent activity of DAF-12 is required for reduced fertility in PD_Stv_ adults. The related gene *nhr-8* is also up-regulated 2.5-fold in PD_Stv_ adults (Lindblom et al. 2001; Magner et al. 2013; Ow et al. 2018); however, *nhr-8(ok186)* exhibited a significant brood size reduction between control and PD_Stv_ animals at levels comparable to wild-type N2 (Fig. 1B; Supplemental Table S1). These results indicate that the reduced fertility of PD_Stv_ adults is dependent on DA biosynthesis through DAF-9 and DAF-36 and the subsequent DA-binding activity of DAF-12.

We next asked whether the exogenous addition of DA was sufficient to reduce the brood size of PD_Stv_ adults. To test this hypothesis, we induced larva into dauer by starvation with or without exogenously added 40 nM of Δ^7^-DA and measured their brood size. The addition of Δ^7^-DA did not affect the fertility of wild-type and *daf-36(k114)* (Fig. 1C; Supplemental Table S2). However, *daf-9(rh50)* PD_Stv_ adults exhibited a significant decrease in brood size in the presence of exogenous Δ^7^-DA (Fig. 1C; Supplemental Table S2). DAF-9 is reported to promote a feedback loop of DA production between the neuroendocrine XXX cells and the hypodermis; thus, *daf-9* mutants may be more sensitive to small changes in DA concentration (Antebi 2014). Taken together, these results suggest that increased DA-dependent DAF-12 signaling is necessary, but perhaps not sufficient, for the reduced brood size phenotype of PD_Stv_ adults.

### The DAF-16 reproductive longevity pathway mediates reproductive plasticity

DAF-16 acts as a major effector of reduced insulin/IGF-1-like signaling (IIS) by promoting the expression of a group of stress response genes (Class I targets) (Tepper et al. 2013). PQM-1, a C2H2 zinc finger transcription factor, acts in a mutually antagonistic manner to DAF-16 by up-regulating genes associated with growth and reproduction (Class II targets), as its cellular localization in the nucleus is anti-correlated to that of DAF-16 (Fig. 2A) (Tepper et al. 2013). Interestingly, we found that the set of genes with significant changes in mRNA levels between PD_stv_ and controls was enriched for DAF-16 Class I and Class II targets (Supplemental Results, Fig. S2A). In addition, we found two genes that regulate DAF-16 cellular localization, *pqm-1* and *daf-18*, were significantly up- and down-regulated, respectively, in PD_stv_ adults compared to controls (Ow et al. 2018). DAF-18 the functional ortholog of the human PTEN tumor suppressor gene that promotes the nuclear localization of DAF-16 (Ogg and Ruvkun, 1998; Gil et al. 1999; Mihaylova et al. 1999; Solari et al. 2005). Together, the overlap in DAF-16 gene targets and the significant changes in the expression of *pqm-1* and *daf-18* prompted us to ask whether PQM-1 and DAF-18 contribute to the reduced fertility in PD_Stv_ adults by altering the regulation of genes involved in reproduction and sequestering DAF-16 in the cytoplasm. We investigated this hypothesis by performing brood assays on adults carrying mutations in *pqm-1* and *daf-18*. Surprisingly, the brood sizes of control and PD_Stv_ adults in *pqm-1(ok485)* and *daf-18(e1375)* strains were similar to wild type, indicating that gene regulation by PQM-1 is unlikely to contribute to the PD_Stv_ decreased brood size phenotype (Fig. 2B; Supplemental Table S4). Since *daf-18(e1375)* is a hypomorph, we next tested the possibility that DAF-16 localization may play a role in regulating postdauer reproduction. *daf-16* null mutants are dauer defective; thus, we asked whether a *daf-16(mu86)* null mutant expressing a rescuing transgene (*daf-16a^AM^::gfp*) that constitutively localizes DAF-16 to the nucleus would show a brood size plasticity phenotype (Lin et al. 2001). Similar to what was observed for *daf-18(e1375)*, the *daf-16(mu86); daf-16a^AM^::gfp* transgenic strain displayed a reduced brood size in PD_Stv_ similar to wild-type N2 (Fig. 2B; Supplemental Table S4). These results suggest that the nuclear localization of DAF-16, and not PQM-1, may be contributing to the diminished fertility phenotype of PD_Stv_ adults.

**Figure 2.**
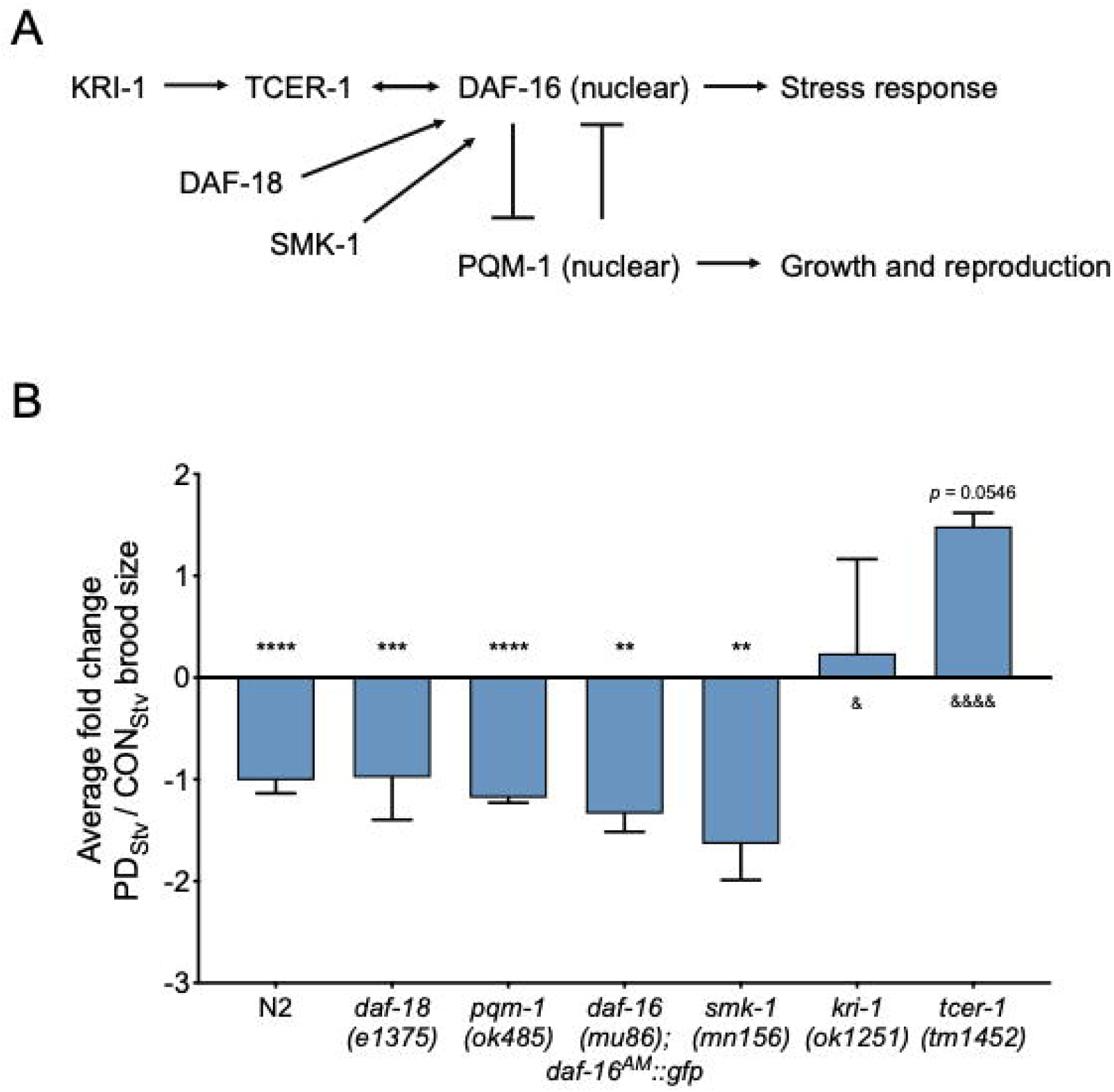
TCER-1 and KRI-1 regulate decreased fertility phenotype in PD_stv_ adults. (*A*) Regulation of DAF-16 nuclear localization. See text for details. (*B*) Brood size comparison of PD_Stv_ relative to CON_Stv_ in wild-type N2 and mutant strains represented as the mean ± S.E.M. ** *p* < 0.01, *** *p* < 0.001, **** *p* < 0.0001; Student’s *t*-test; comparison of PD_Stv_ and CON_Stv_ of a given genotype. ^&^ *p* < 0.05, ^&&&&^ *p* < 0.0001; one-way ANOVA with Fisher’s LSD *post hoc* test; comparison of the PD_Stv_/CON_Stv_ brood size change between a given mutant genotype to wild-type N2. Additional brood size data is provided in Supplemental Table S4.

To further investigate whether the nuclear localization of DAF-16 is required for the reproductive plasticity between control and PD_Stv_ animals, we measured the brood size of animals with mutations in genes that regulate the cellular localization of DAF-16. First, we tested SMK-1/PPP4R3A, which promotes the nuclear localization of DAF-16 when animals are exposed to pathogenic bacteria, ultraviolet irradiation, and oxidative stress, but does not affect the dauer development and reproductive timing activities of DAF-16 (Wolff et al. 2006). We found that *smk-1(mn156)* mutants continued to exhibit a decreased PD_Stv_ fertility compared to controls. Secondly, we examined two genes, *kri-1* (an ortholog of the human intestinal ankyrin-repeat protein KRIT1/CCM1) and *tcer-1* (a homolog of the human transcription elongation factor TCERG1), that are required for DAF-16 nuclear localization and extend adult lifespan in germ line-less animals, but do not regulate dauer formation (Berman and Kenyon, 2006; Ghazi et al. 2009). We found that decreased brood size in PD_Stv_ was abrogated in the *kri-1(ok1251)* and *tcer-1(tm1452)* mutants (Fig. 2B; Supplemental Table S4). Because TCER-1 regulates the transcription of its target genes in a DAF-16-dependent and independent manner (Armit et al. 2016), these results indicate that KRI-1, TCER-1, and possibly DAF-16 are required for the reproductive plasticity in PD_Stv_ animals.

### Increased fatty acid metabolism promotes PD_Stv_ fertility

In animals lacking a germ line, DAF-16 and TCER-1 are required to bolster the expression of lipid biosynthesis, storage, and hydrolysis genes to promote adult longevity (Armit et al. 2016). Similar to germ line-less *glp-1* mutants, PD_Stv_ adults exhibited a significant altered expression of ~26% (33 of 127 genes) of all the fatty acid metabolic genes, including ~46% (18 of 39 genes) also targeted by DAF-16 and TCER-1 (Fig. S2B; Supplemental Table S5). These results suggest that DAF-16 and TCER-1 play a role in regulating reproductive plasticity between controls and PD_Stv_ by up-regulating fatty acid metabolism.

To investigate this hypothesis, we asked if mutations in known regulators of lipid metabolism would exhibit changes in brood size in PD_Stv_ adults when compared to controls. One of the genes jointly up-regulated by DAF-16 and TCER-1 is *nhr-49*, a nuclear hormone receptor homologous to the mammalian HNF4α lipid sensing nuclear receptor involved in the regulation of fatty acid metabolism and the oxidative stress response (Ratnappan et al. 2014; Amrit et al. 2016; Moreno-Arriola et al. 2016; Goh et al. 2018; Hu et al. 2018). Additional nuclear hormone receptors, NHR-80, NHR-13, and NHR-66, and the Mediator complex subunit, MDT-15, partner with NHR-49 and co-regulate the expression of genes involved in various aspects of lipid metabolism such as fatty acid β-oxidation, transport, remodeling, and desaturation (Van Gilst et al. 2005a, 2005b; Taubert et al. 2006; Nomura et al. 2010; Pathare et al. 2012; Ratnappan et al. 2014; Folick et al. 2015; Amrit et al. 2016). In addition, SBP-1 (homolog of mammalian SREBP) and NHR-49 are co-regulated by MDT-15 as part of a transcriptional network coordinating the expression of delta-9 (Δ9) fatty acid desaturase genes (Fig. 3B) (Yang et al. 2006; Taubert et al. 2006).

**Figure 3.**
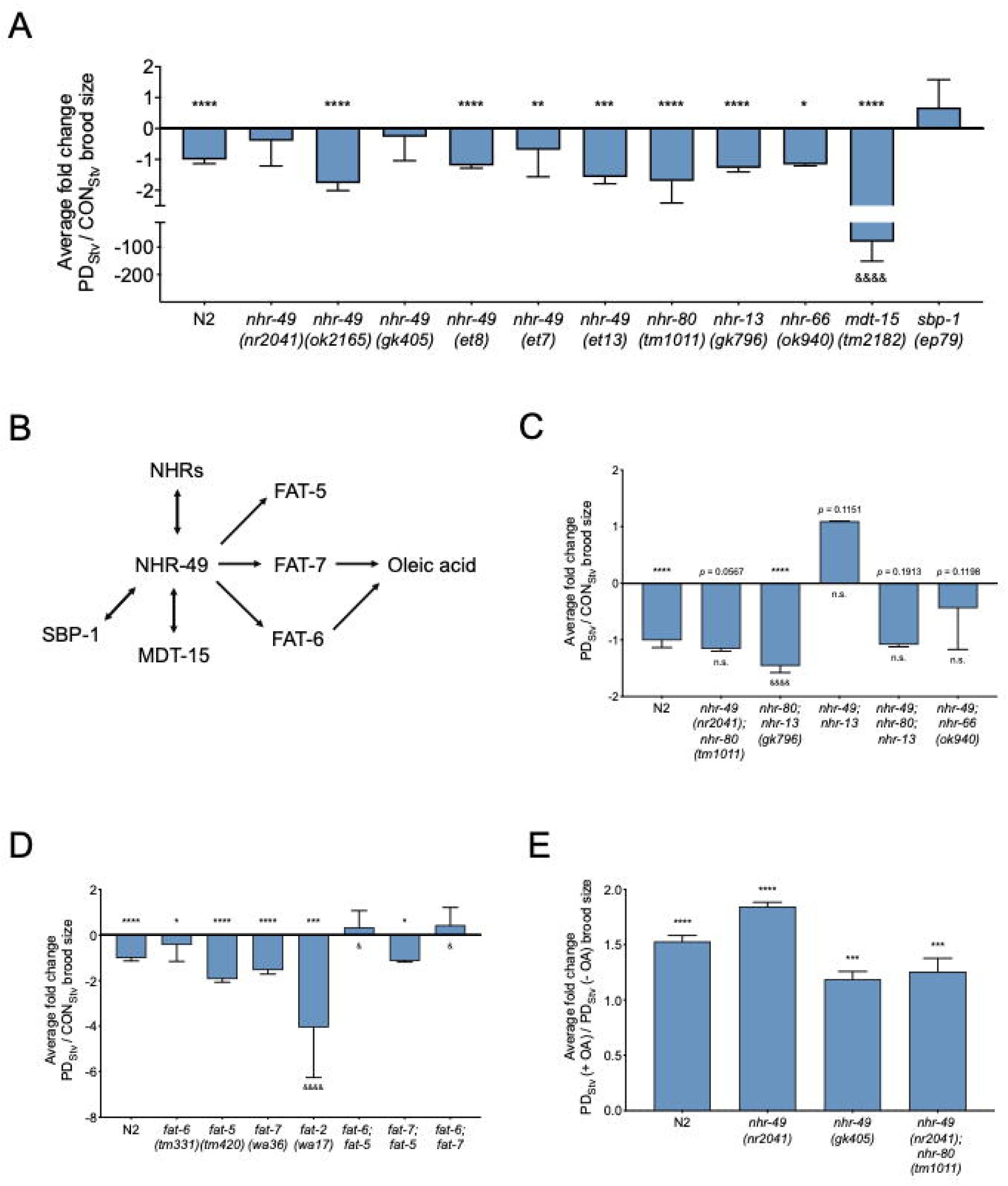
NHR-49, SBP-1, and FAT-6 modulate adult reproductive plasticity. (*A,C,D*) Brood size comparison of PD_Stv_ relative to CON_Stv_ for wild-type N2 and mutant strains represented as the mean ± S.E.M. Additional brood size data is provided in Supplemental Tables S6, S7, and S8. (*B*) Model of lipid metabolism regulators. See text for details. (*E*) Brood size of PD_Stv_ wild-type N2 and mutant animals fed with *E. coli* OP50 pre-loaded with or without 600 μM of oleic acid (OA), represented as the mean ± S.E.M. Additional brood size data is provided in Supplemental Table S9. * *p* < 0.05, ** *p* < 0.01, *** *p* < 0.001, **** *p* < 0.0001; Student’s *t*-test; comparison of PD_stv_ and CON_stv_ brood size within a given genotype. ^&^ *p* < 0.05, ^&&&&^ *p* < 0.0001 and n.s. (no significance); one-way ANOVA with Fisher’s LSD post hoc test; comparison of PD_Stv_/CON_Stv_ brood size between a given genotype to wild-type N2.

The reduced fertility of PD_Stv_ characteristic of wild-type animals was also observed in *nhr-80(tm1011)*, *nhr-13(gk796)*, *nhr-66(ok940)*, and *mdt-15(tm2182)* strains, but not in *sbp-1(ep79)* (Fig. 3A; Supplemental Table S6). In addition, two *nhr-49 loss-of-function* (*lof*) mutants, *nhr-49(nr2014)* and *nhr-49(gk405)*, failed to exhibit the decreased fertility in PD_Stv_ adults compared to CON_Stv_, while a third *nhr-49(ok2165) lof* allele continued to display a PD_Stv_ reduction in brood size (Fig. 3A; Supplemental Table S6). Because the three *nhr-49* mutant alleles, *nr2014*, *gk405*, and *ok2165*, differ in the nature and the location of their lesions (Lee et al. 2016), their biological function may vary and hence result in phenotypic differences (*nr2041* and *gk405* vs. *ok2165*). Animals expressing *nhr-49* gain-of-function alleles (*et7*, *et8*, and *et13*) with missense lesions located at or near the LBD may augment NHR-49 activity resulting in a reproductive plasticity phenotype comparable to that of wild-type N2 (Fig. 3A; Supplemental Table S6) (Svensk et al. 2013; Lee et al. 2016). Because of the interaction between NHR-49, NHR-80, NHR-13, and NHR-66, we also examined if double and triple mutants of these NHRs would have any reproductive plasticity phenotypes. Double and triple mutant combinations of alleles exhibited an abolishment of the PD_Stv_ decreased brood size phenotype only if they expressed the *nhr-49(nr2041)* mutation (Fig. 3C; Supplemental Table S7). Thus far, our results indicate that NHR-49 and SBP-1 are important in the reproduction program of PD_Stv_ adults, likely by up-regulating fat metabolism genes.

NHR-49 and SBP-1 up-regulate the expression of genes involved in fatty acid biosynthesis, including the Δ9 desaturases, *fat-5*, *fat-6*, and *fat-7*, and the delta-12 (Δ12) desaturase *fat-2* (Nomura et al. 2010; Han et al. 2017). FAT-5, FAT-6, and FAT-7 convert saturated fatty acids (SFAs) to mono-unsaturated fatty acids (MUFAs), while FAT-2 catalyzes the conversion of MUFAs to poly-unsaturated fatty acids (PUFAs) (Watts and Ristow, 2017). Our previous mRNA-Seq results showed that the expression of *fat-5*, *fat-6*, *fat-7*, and *fat-2* increased significantly between 3.8- and 26.6-fold in wild-type PD_Stv_ adults compared to controls (Ow et al. 2018). While strains expressing a single *fat-5(tm331)*, *fat-6(tm420)*, *fat-7(wa36)*, or *fat-2(wa17)* mutation continued to exhibit the reduced brood size phenotype in PD_Stv_ adults, double mutants expressing a mutation in *fat-6* no longer displayed the reduced PD_Stv_ phenotype. The double mutant *fat-7(wa36); fat-5(tm331)* continued to display a significant fertility difference between controls and PD_Stv_ (Fig. 3D; Supplemental Table S8). These results suggest: 1) a functional redundancy between the Δ9 fatty acid desaturases in modulating lipid homeostasis of PD_Stv_ adults, with FAT-6 playing a more principal role than FAT-5 and FAT-7; and 2) MUFAs are required for the decreased fertility phenotype in PD_Stv_ adults compared to controls.

In *C. elegans*, MUFAs are essential for viability as a triple *fat-5; fat-6; fat-7* mutant is lethal (Brock et al. 2006). MUFAs, such as oleic acid, can be remodeled to become PUFAs, phospholipids, and neutral lipids such as triacylglycerols (TAGs), which serve as energy storage molecules in the intestine, hypodermis, and germ line (Watts and Ristow, 2017). In addition to acting as key regulators of fat metabolism, FAT-5, FAT-6, and FAT-7 are also essential in promoting the long lifespan of adult worms lacking a germ line (Van Gilst et al. 2005a; Brock et al. 2006; Goudeau et al. 2011; Ratnappan et al. 2014). Given that MUFAs are required for the reduced fertility of PD_Stv_ adults compared to controls, we asked whether the dietary addition of oleic acid to PD_Stv_ animals would further reduce their brood size such that they would produce even fewer progeny compared to PD_Stv_ animals not supplemented with oleic acid. To test this hypothesis, we compared the brood size of PD_Stv_ adults fed *E. coli* OP50 grown with oleic acid and PD_Stv_ adults whose bacterial diet was not pre-loaded with oleic acid. We tested the N2 wild-type, *nhr-49(nr2041)*, *nhr-49(gk405)*, and *nhr-49(nr2041); nhr-80(tm1011)* strains, and found that for all genotypes, the supplementation of oleic acid significantly increased the progeny number produced by PD_Stv_ compared to PD_Stv_ adults on the control diet (Fig. 3E; Supplemental Table S9). These results indicate that oleic acid is a limiting factor for reproduction, even for wild-type animals, after passage through the starvation-induced dauer stage. Altogether, these results suggest that NHR-49 and SBP-1 are critical for the decreased fertility in PD_Stv_ adults by mediating the up-regulation of fatty acid desaturases and promoting the synthesis of sufficient levels of oleic acid needed for reproduction after the animals experienced starvation-induced dauer formation.

### Starvation-induced postdauer adults have reduced fat stores

In long-lived *glp-1* mutants lacking a germ line, fat stores are increased relative to wild-type (Steinbaugh et al. 2015; Amrit et al. 2016). Given that PD_Stv_ adults also have an increased expression of lipid metabolism genes similar to *glp-1* adults, but have limited quantities of oleic acid for reproduction, we questioned what the status of fat stores would be in wild-type adults with an intact germ line that had passed through dauer due to starvation. Using Oil Red O (ORO) staining, we compared the amounts of neutral triglycerides and lipids (O’Rourke et al. 2009) in PD_Stv_ one-day-old adults and age-matched control adults. Interestingly, PD_Stv_ adults have a significantly reduced amount of stored lipids relative to control adults in their intestine, but still displayed ORO staining in the embryos within the uterus (Fig. 4A, 4B). These results are consistent with a model that PD_Stv_ adults have increased expression of lipid metabolism genes in order to produce sufficient levels of MUFAs needed for reproduction rather than lipid storage.

**Figure 4.**
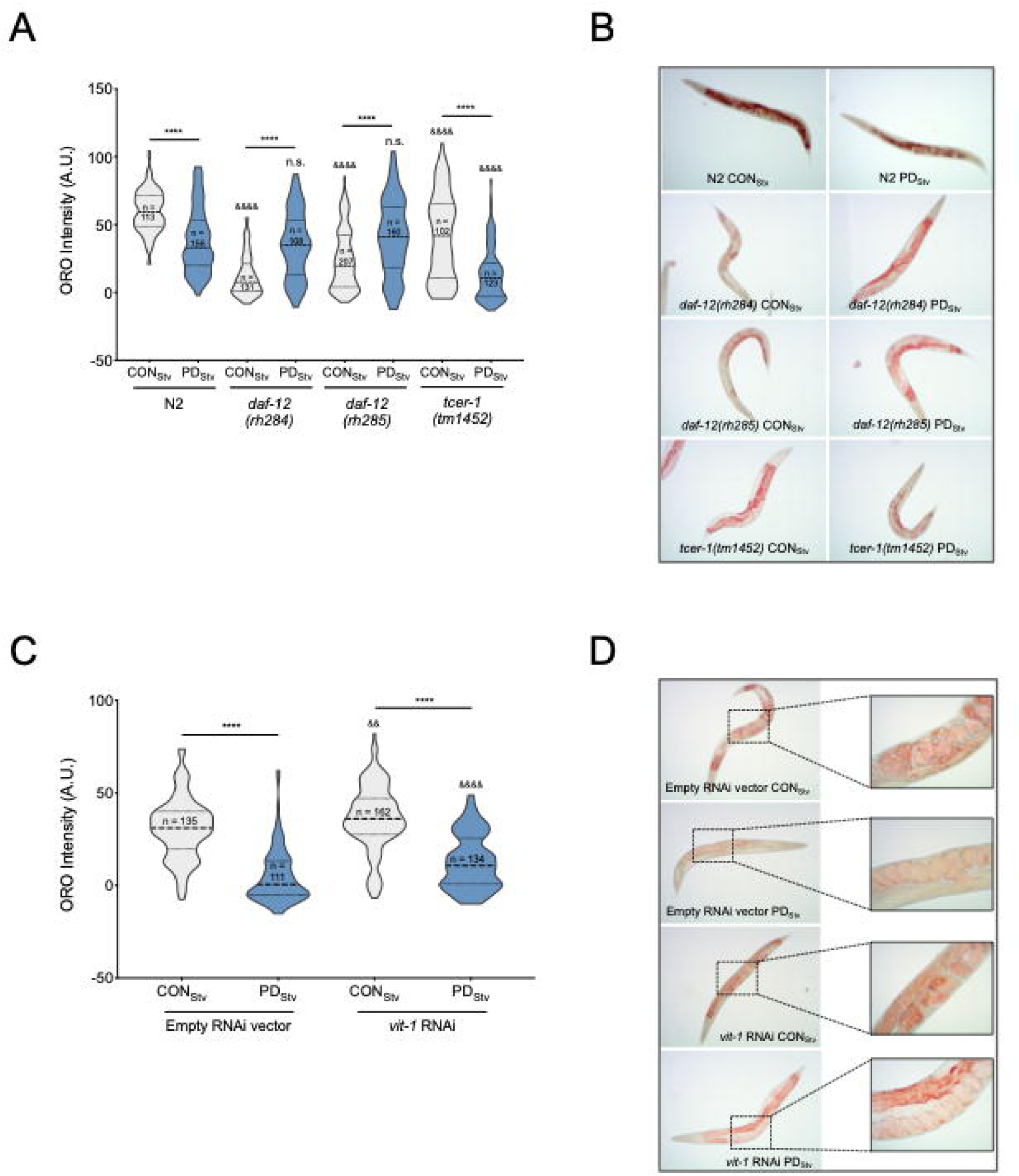
Fat stores are reduced in wild-type postdauer adults that experienced starvation-induced dauer. (*A, C*) Oil Red O (ORO) intensity in CON_Stv_ and PD_Stv_ adults, represented as violin plots with the mean (dashed line), first, and third quartiles indicated (dotted lines). (*A*) **** *p* < 0.0001; Student’s *t*-test; comparison between CON_Stv_ and PD_Stv_ of the same genotype. ^&&&&^ *p* < 0.0001; one-way ANOVA with Fisher’s LSD *post hoc* test comparing mutant CON_stv_ or PD_stv_ to wild-type N2. n.s.; no significance. Total sample size is indicated by *n*. (*C*) **** *p* < 0.0001; Student’s *t*-test; comparison between CON_Stv_ and PD_Stv_ of the same condition. ^&&^ *p* = 0.0038 and ^&&&&^ *p* < 0.0001; Student’s *t*-test; comparison of CON_stv_ or PD_stv_ *vit-1* RNAi to empty RNAi vector control. (*B, D*) Representative micrographs of one-day-old adults stained with ORO.

Next, we investigated whether the decreased lipid stores in PD_Stv_ adults was dependent upon DAF-12 and TCER-1. Both *daf-12(rh284)* and *daf-12(rh285)* mutants displayed a significant increase in lipid storage in PD_Stv_ adults relative to control adults, opposite to what was observed in wild-type, indicating that DA-mediated DAF-12 activity is required for regulation of fat stores in PD_Stv_ adults (Fig. 4A, 4B). Interestingly, the ORO staining positively correlated with brood sizes in wild-type N2 and the *daf-12* mutants: fat stores and fertility decreased in wild-type PD_Stv_ adults, while fat stores and fertility increased in *daf-12* PD_Stv_ adults compared to *daf-12* CON_Stv_ adults (Fig. 1B, 4A, 4B; Supplemental Table S1). In *tcer-1(tm1452*) adults, lipid staining was diminished in PD_Stv_ compared to controls, similar to what is seen in wild-type (Fig. 4A, 4B). However, since both *tcer-1* control and PD_Stv_ adults have reduced fat stores compared to their wild-type counterparts, and TCER-1 is known to positively regulate NHR-49 and fatty acid metabolism, this result is likely due to the inability of these animals to store fat and not because the decreased fat stores in wild-type PD_Stv_ adults is TCER-1 independent. In addition to regulation of fat metabolism genes in the intestine, TCER-1 also functions redundantly with PUF-8, a member of an evolutionarily conserved stem cell proliferation modulatory family, to potentiate germ cell proliferation in animals undergoing continuous development (Pushpa et al. 2013). Thus, our *tcer-1(tm1452)* results suggest that TCER-1 activity may also be necessary in regulating germline mechanisms that curtail fertility following exit from dauer diapause.

During *C. elegans* reproduction, intestinal fat stores are reallocated into low-density lipoproteins (LDL)-like particles (yolk lipoproteins or vitellogenins) that are incorporated into oocytes through receptor-mediated endocytosis in a process called vitellogenesis, and serve as the only supply of nutrients to the developing embryos (Kimble and Sharrock, 1983; Grant and Hirsch, 1999). Six vitellogenins homologous to the human ApoB proteins are encoded in the *C. elegans* genome, *vit-1* through *vit-6*, and concomitant multiple RNAi knockdown of the *vit* genes increases adult lifespan in a process that requires NHR-49 and NHR-80 (Spieth et al. 1991; Seah et al. 2016). The transition from dauer to reproductive adulthood is an energetically costly process that likely drains the already reduced fat reservoirs in dauers that experienced early-life starvation. Because vitellogenesis mobilizes intestinal fat resources for reproduction and depletes somatic lipid stores (Kimble and Sharrock, 1983), we hypothesized that PD_Stv_ adults have reduced fat reservoirs because they prioritize vitellogenesis as a reproductive investment over intestinal storage. To test this hypothesis, we examined the fat stores in control and PD_Stv_ adults following RNAi knockdown of *vit-1*, which also results in the knockdown of *vit-2/3/4/5* due to the high sequence homology amongst the *vit* genes (Seah et al. 2016) (Supplemental Fig. S4). In the RNAi treated animals, PD_Stv_ adults continued to exhibit decreased fat stores compared to control adults, similar to the empty vector negative controls. However, PD_Stv_ adults treated with *vit* RNAi have significantly greater intestinal fat deposits than PD_Stv_ negative controls (*p* < 0.0001), indicating that increased vitellogenesis in PD_Stv_ adults may be one contributing factor to the lack of stored intestinal fat (Fig. 4C, 4D). These results are consistent with our previous finding that oleic acid is a limiting resource for reproduction in PD_Stv_ animals, and suggests a model where PD_Stv_ adults utilize fat accumulated after diapause for reproduction and not storage.

### Steroid signaling, reproductive longevity, and fatty acid metabolic pathways act synergistically at different developmental time points to regulate reproductive plasticity

Thus far, our results indicate that both DA-dependent DAF-12 activity and TCER-1 are required for the reduced fertility phenotype in PD_Stv_ adults. However, the *daf-12* and *tcer-1* PD_Stv_ mutant phenotypes are distinct, suggesting that they act in different pathways, tissues, or developmental time points to regulate PD_Stv_ fertility. Furthermore, the brood sizes of *kri-1(ok1251)* or *tcer-1(tm1425)* PD_Stv_ adults supplemented with exogenous Δ^7^-DA were unaffected, indicating that the effect of DA on PD_Stv_ brood size is independent of the reproductive longevity pathway (Supplemental Fig. S3; Supplemental Table S12). To further investigate the developmental mechanism of steroid signaling, reproductive longevity, and fatty acid metabolism pathways in the regulation of reproductive plasticity, we examined if DAF-12, TCER-1, and NHR-49 play a direct role in the timing of germline proliferation in postdauer larvae. We previously demonstrated that wild-type PD_Stv_ animals delay the onset of germline proliferation compared to control animals, resulting in a reduced brood size (Ow et al. 2018). *C. elegans* begins germ cell proliferation after hatching and continues throughout larval development. Germ cells at the most distal ends of the gonad undergo mitotic self-renewal while germ cells located more proximal in the gonad enter and undergo meiosis (Kimble and Crittenden, 2005). In *C. elegans* hermaphrodites, undifferentiated germ cells initiate spermatogenesis during the L4 larval stage, followed by a transition to oogenesis at the adult stage (L’Hernault, 2006). Therefore, reproduction in *C. elegans* hermaphrodites is sperm-limited (Byerly et al. 1976; Ward and Carrel, 1979; Kimble and Ward, 1988). In previous results, when control and PD_Stv_ somatic development was synchronized using vulva morphology, we observed significantly fewer germ cell rows in PD_Stv_ larvae compared to control larvae, correlating with fewer sperm available for self-fertilization in adulthood (Ow et al. 2018).

To determine if DAF-12, TCER-1, and NHR-49 play a direct role in germline proliferation as a mechanism to regulate reproductive plasticity, we counted the number of germ cell rows in control and PD_Stv_ mutant larvae that were developmentally synchronized by their somatic morphology. Because *daf-12(rh284)* and *daf-12(rh285)* mutants have altered germline morphologies that prevent accurate synchronization, we used the *daf-36(k114)* mutation to disrupt the steroid signaling pathway. First, we recapitulated our previous results showing that wild-type PD_Stv_ larvae have significantly fewer total germ cell rows compared to control larvae due to reduced cell rows in the meiotic transition zone (Fig. 5; Supplemental Fig. S6). In contrast, *daf-36(k114)* control and PD_Stv_ larvae did not exhibit a significant difference in total (mitotic and meiotic) cell rows, indicating that DAF-36-dependent DA is required during early germline development for the delay in PD_Stv_ germ cell proliferation (Fig. 5; Supplemental Fig. S6). This result is consistent with the increased brood size of PD_Stv_ adults in *daf-12(rh284)* and *daf-12(rh285)* mutants, which express DAF-12 proteins unable to bind to DA.

**Figure 5.**
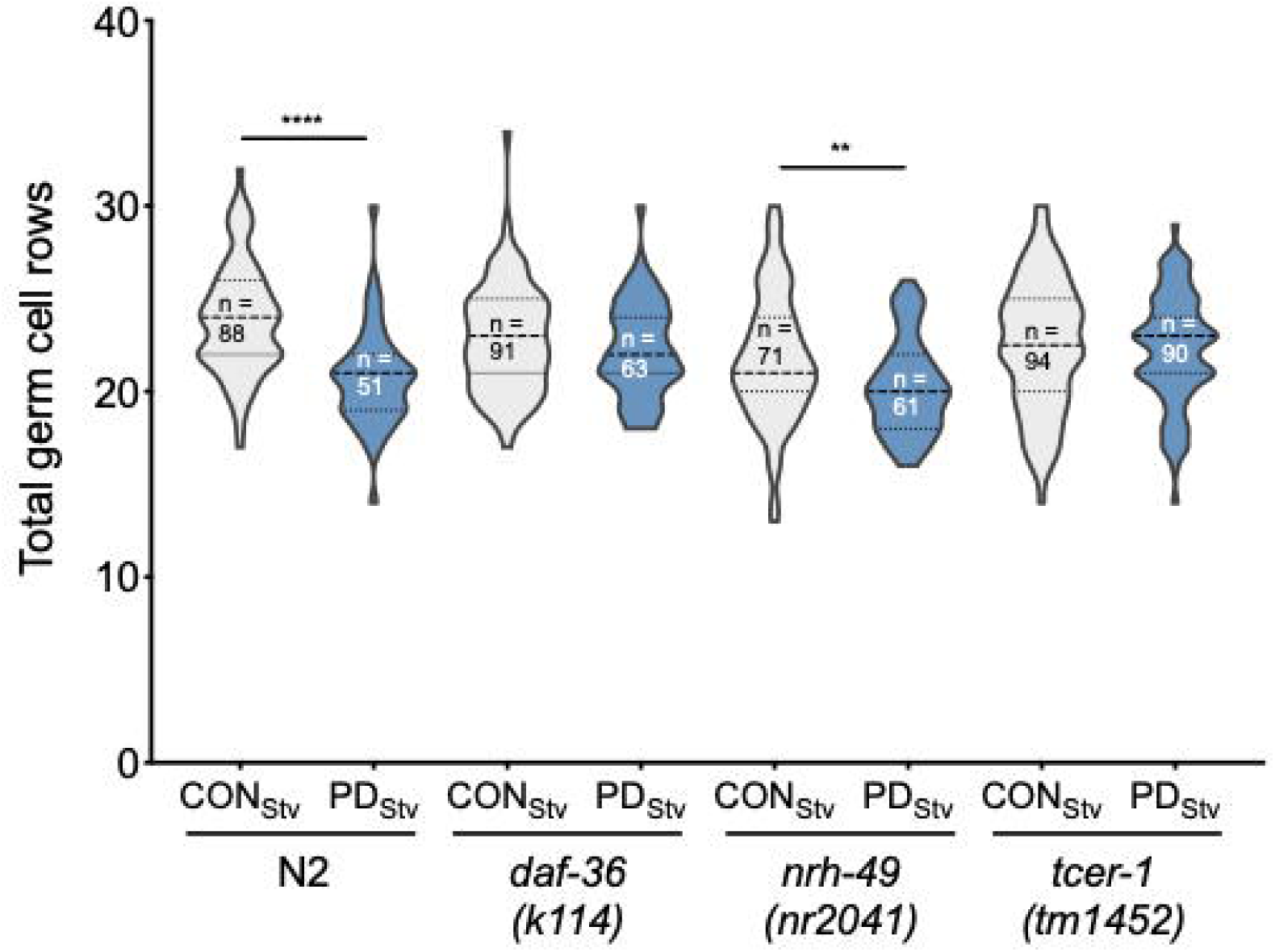
DAF-12 and TCER-1 regulate the onset of germline proliferation. Total germ cell rows in CON_Stv_ and PD_Stv_ wild-type N2 and mutant larva exhibiting L3 vulva morphology (see Materials and methods). Data represented as violin plots with the mean (dashed line), first, and third quartiles indicated (dotted lines) ** *p* < 0.01; **** *p* < 0.0001; Student’s *t*-test. Total sample size is indicated by *n*.

In contrast to the *daf-36* mutant, the *nhr-49(nr2041)* mutant exhibited a significant decrease in the number of total germ cell rows in PD_Stv_ larvae relative to control larvae, similar to wild-type animals (Fig. 5). However, the numbers of germ cell rows in the mitotic and meiotic zones of *nhr-49* control and PD_stv_ animals were not significantly different (Supplemental Fig. S6). Since NHR-49 is required for the reduced fertility in PD_Stv_ adults, these results suggest that NHR-49 plays a small role in regulating germline proliferation in larvae, and a larger role in regulating fatty acid metabolism in the intestine to support vitellogenesis in adults. Like NHR-49, TCER-1 plays a role in the intestine acting to upregulate fatty acid metabolism genes in animals lacking germ cells, but also plays a role in the regulation of germ cell proliferation (Pushpa et al. 2013; Amrit et al. 2016). We found that a mutation in *tcer-1* resulted in a similar number of total germ cell rows in control and PD_Stv_ larvae, indicating that TCER-1 plays a role in PD_Stv_ reproductive plasticity by regulating germ cell proliferation in addition to its intestinal role (Fig. 5). Interestingly, *tcer-1* PD_Stv_ larvae also appear to have a defect in the onset of meiosis, since PD_Stv_ larvae have significantly great number of mitotic germ cell rows compared to controls (Supplemental Fig. S6). Together, these results indicate that DAF-12 steroid signaling and TCER-1 act early in germline development to delay the onset of germ cell proliferation in PD_Stv_ animals, while NHR-49 primarily functions later in development to promote the reduced fertility of PD_Stv_ adults.

### Generational transmission of early life starvation memory

Our results suggest that PD_Stv_ upregulate their fatty acid metabolism to increase fat transport to embryos. In humans, nutritional stress *in utero* not only promotes metabolic syndrome in adulthood of the affected individuals, but also promotes obesity in subsequent generations (Painter et al. 2008; Veenendaal et al. 2013). Therefore, to examine whether the starvation-induced physiology of PD_Stv_ adults is inherited, we used ORO staining to quantitate the fat storage in adult F_1_ progeny of control and PD_Stv_ adults that did not experience starvation themselves. Indeed, the adult F_1_ progeny of PD_Stv_ adults had an increased level of stored fat compared to F_1_ progeny of control adults (Fig. 6), but the difference was abolished by the F_2_ generation. These results indicate that the F_1_ progeny of PD_Stv_ adults inherit a starvation memory that results in metabolic reprogramming to increase their stored fat reserves.

**Figure 6.**
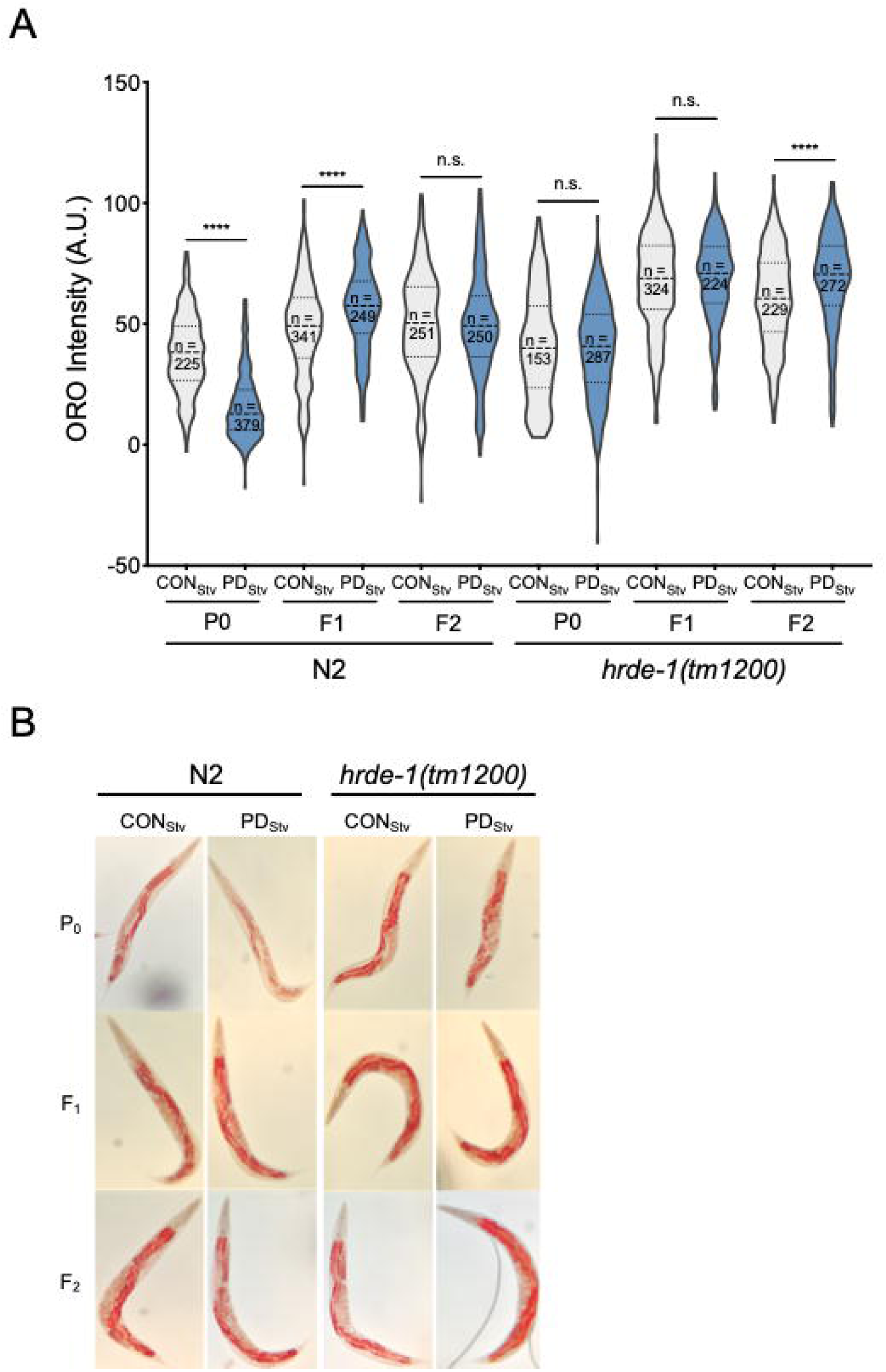
Generational inheritance of starvation memory. *(A)* Intensity of ORO lipid staining of wild-type N2 and *hrde-1(tm1200)* control and postdauer P_0_, F_1_, and F_2_ generations, represented as violin plots with the mean (dashed line), first, and third quartiles indicated (dotted lines). **** *p* < 0.0001; Student’s *t*-test. n.s.; no significance. Total sample size is indicated by *n*. (*B*) Representative micrographs of N2 and *hrde-1(tm1200)* CON_Stv_ and PD_Stv_ P_0_, F_1_, and F_2_ worms stained with ORO.

In *C. elegans*, inheritance of gene expression states is often mediated by small RNA pathways (Feng and Guang, 2013; Rechavi and Lev, 2017). The germline-specific, nuclear Argonaute HRDE-1/WAGO-9 has been shown to promote transgenerational silencing via the formation of heterochromatin at targeted genomic loci (Buckley et al. 2012; Feng and Guang, 2013; Rechavi and Lev, 2017). In particular, starvation-induced L1 diapause has been shown to alter the small RNA populations of subsequent generations in a HRDE-1 dependent manner (Rechavi et al. 2014). We examined whether HRDE-1 was required for the generational inheritance of starvation memory by quantifying fat storage in *hrde-1(tm1200)* mutant control and PD_Stv_ adults and their F_1_ and F_2_ progeny. First, we observed that the stored fat levels between *hrde-1* control and PD_Stv_ adults were similar, suggesting that the germline-specific Argonaute is required for decreased intestinal fat storage of PD_Stv_ adults (Fig. 6). In addition, the ORO staining of the F_1_ progeny of control and PD_Stv_ adults was also similar, indicating that HRDE-1 is required for the generational inheritance of starvation memory (Fig. 6). Remarkably, in the F_2_ generation, the grand-progeny of *hrde-*1 PD_Stv_ mutants displayed increased ORO staining like that observed for the F_1_ generation of wild-type adults, suggestive of a generational delay in the physiological response to ancestral early-life starvation upon the loss of HRDE-1 activity (Fig. 6). Our genetic evidence suggests that the germline nuclear RNAi pathway plays two roles: it contributes to the metabolic programming of individuals that experienced early life starvation as well as promotes the generational inheritance of starvation memory in the progeny. Our results show that *C. elegans*, like humans, inherit the disposition for increased adiposity from parents that experienced early life starvation.

## Discussion

The trade-off between reproduction and longevity has long been associated with the notion that in the absence of reproduction, the fat stores of an animal would be redistributed to promote somatic maintenance (Williams 1966; Kirkwood 1977; Westendorp and Kirkwood, 1998; Partridge et al. 2005). As such, reducing or suppressing reproduction in animals often extends lifespan, in part due to enhanced lipogenic processes (Fowler and Partridge 1989; Kenyon 2010, 2010b; Judd et al. 2011). What remains unclear is how reproduction and somatic maintenance are balanced by developmental signals. In this study, we show that starvation early in the life of a *C. elegans* animal with an intact reproductive system programs its transcriptome to promote reproduction. Because dauers are non-feeding, starvation-induced dauers have limited stored fat to survive the diapause stage and remodel out of dauer. We propose a model whereby steroid signaling, reproductive longevity, and fatty acid metabolic pathways are reprogrammed in animals that experienced starvation-induced dauer in order to delay the onset of germline proliferation and redistribute intestinal fat to developing oocytes (Fig. 7). These changes allow PD_Stv_ adults to delay reproduction until energy thresholds are met to provide adequate levels of nutrition to fewer albeit viable embryos at the expense of somatic survival and maintenance. In the absence of a functional germ line, the fat that would normally be allocated to the progeny is channeled to nurture somatic tissues and, consequently, extend adult lifespan. Thus, the mechanisms that prolong lifespan in the absence of a functional germ line are the same cellular programs deployed to respond to developmental signals triggered by starvation early in life history in an animal capable of reproduction.

**Figure 7.**
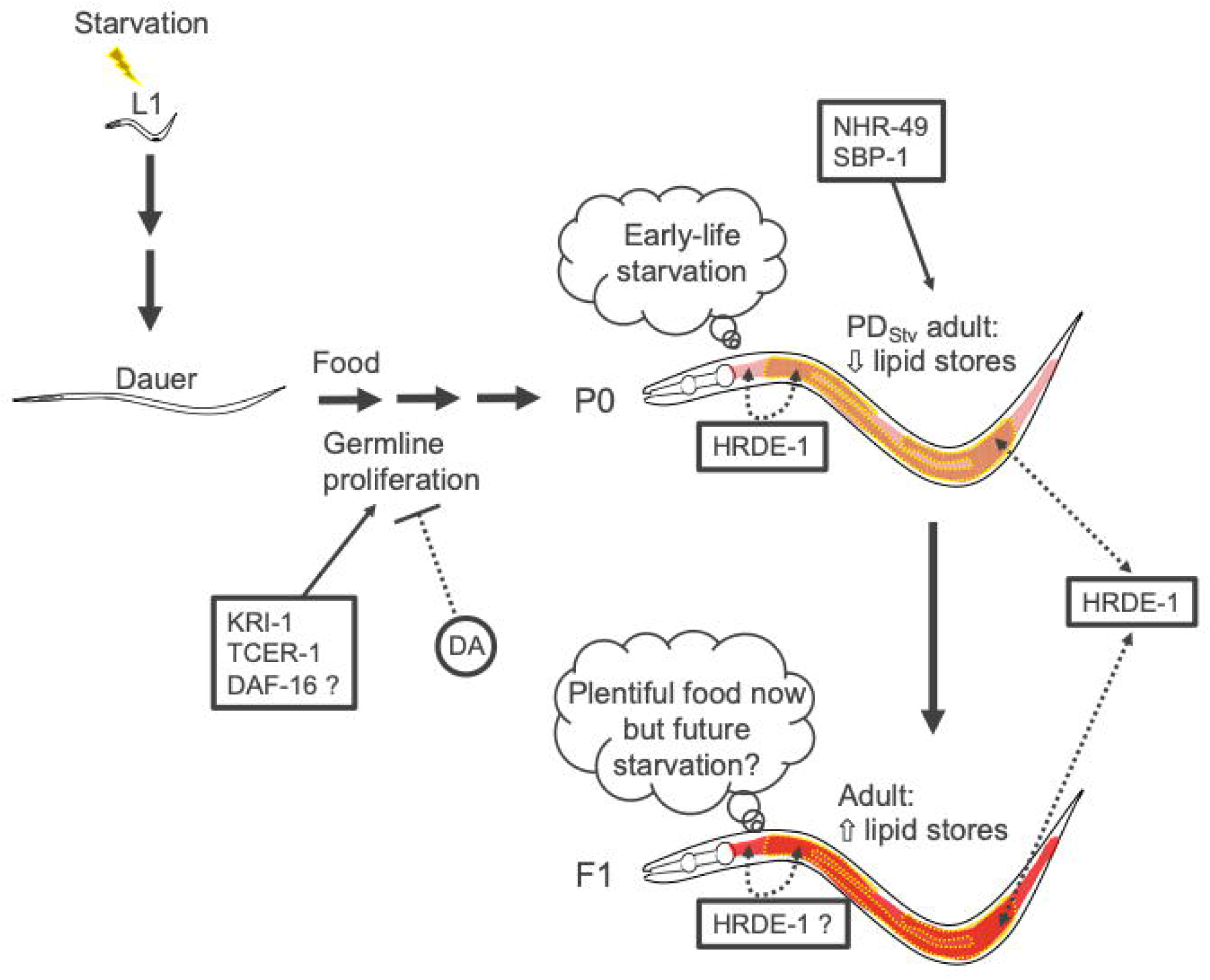
Model depicting endocrine signaling pathways acting as contributors to fatty acid metabolism of PD_stv_ animals. Signaling between the soma and the germ line establishing metabolic plasticity and the transmission of ancestral starvation memory are dependent on HRDE-1.

The extended lifespan associated with animals missing a functional germ line is specifically dependent on the lack of proliferating germ cells and not due to sterility resulting from sperm, oocyte, or meiotic precursor cell deficiency (Hsin and Kenyon, 1999; Arantes-Oliveira et al. 2002). Animals without a functional germ line have up-regulated fat metabolism pathways and exhibit increased levels of intestinal fat stores that are associated with a longer lifespan (Kenyon 2010, 2010b; Amrit et al. 2016). Postdauer adults that experienced early life starvation have a delay in the onset of germ cell proliferation after dauer exit and similarly have upregulated fat metabolism pathways in adulthood. These results indicate that the cellular mechanisms that promote germline proliferation during a critical period in development also negatively regulate lipid metabolism, leading to altered somatic fat accumulation and longevity (Wang et al. 2008). While a number of genes and cellular components affecting germline proliferation have been extensively investigated (Kimble and Crittenden, 2005), what might the actual signals communicating the state of proliferating germ cells be that arbitrate lipid levels and the aging process? Because upregulation of DAF-36 and Δ^7^-DA is crucial for aging extension in germline-less animals and reproductive plasticity in PD_Stv_ adults, it is likely that the signal communicating the state of germ cell proliferation may include dafachronic acids. Dafachronic acids mediating increased longevity are produced in the somatic gonad, which includes the stem cell niche site of the germ line (Yamawaki et al. 2010). Based on our results, DA is required for the delay in germline proliferation (Fig. 5), but DA-dependent DAF-12 activity is not necessary for the increase in fatty acid metabolism observed in PD_Stv_ adults (Fig. 4A, 4B), suggesting the possibility that dafachronic acids act in a DAF-12-independent manner to regulate germ cell proliferation and fat metabolism. In addition, our results showed that *hrde-1* is required for the altered fat metabolism in PD_Stv_ adults (Fig. 6), raising the possibility that small RNAs could be another type of signaling molecule communicating the state of the germ line to the soma. Small RNA pathways are potent surveillance mechanisms that guard against the threat of foreign genetic invaders such as viruses and transposons which can compromise germline integrity (Almeida et al. 2019). Because germ line proliferation occurs throughout the lifetime of *C. elegans*, it is likely that small RNA pathways survey the proliferative nature of the germ line at all stages of animal development and potentially act as signaling molecules to regulate somatic processes dependent upon the germ line.

One of the most intriguing findings of this study is that the starvation memory of the parent is “bequeathed” to the F_1_ progeny, triggering elevated levels of fat stores, presumably as a physiological defense against ancestral famine (Fig. 6). Interestingly, the increased intestinal fat inherited by the F_1_ progeny of wild-type postdauer parents bypassed the F_1_ generation in the *hrde-1* mutant and was only evident in the F_2_ generation (Fig. 6). One potential explanation is that small RNA signals are transmitted to the F_1_ generation via the HRDE-1 nuclear RNAi pathway to effect somatic phenotypes. However, with the exception of DAF-16, none of the germline longevity pathway genes or the vitellogenesis genes examined in this study are categorized as direct HRDE-1 targets (Buckley et al. 2012). Given that the life stage (adulthood) at which the HRDE-1 targets were identified is the same life stage that was used in this study, we speculate that HRDE-1 may be indirectly targeting endocrine and vitellogenins genes by: 1) targeting germ line genes that then affect somatic gene expression; or 2) indirectly regulating the function of the endocrine and vitellogenin genes by targeting a different repertoire of somatic targets. Interestingly, we find that 62% of small RNAs associated with HRDE-1 target genes (984 out of 1587) are expressed in somatic tissues, such as neurons, intestine, hypodermis, and muscle (Ortiz et al. 2014; Kaletsky et al. 2016). While silencing somatic genes in the germ line is consistent with the described function of HRDE-1, our results raise the possibility that HRDE-1 may also directly, or indirectly, target somatic genes. Accordingly, HRDE-1 is known to contribute to the heritability of a cohort of small RNAs targeting nutrition and lipid transporter genes that was inherited for at least three generations from populations that experienced L1 larval arrest (Rechavi et al. 2014). In addition, HRDE-1 is required for the repression of a group of genes activated upon multi-generational high temperature stress that is inherited for at least two generations in the absence of the stress (Ni et al. 2016). Thus, it is likely that HRDE-1 and the nuclear RNAi pathway may serve as a signaling referee between the soma and the germ line to effect changes due to environmental and developmental signals to perdure ancestral starvation memory.

Our study showed that PD_Stv_ adults have upregulated expression of lipid metabolism genes as a means to load embryos with increased fat and potentially protect progeny against future starvation conditions. Not only are fatty acids critical for cell and organelle maintenance and efficient energy storage molecules but they can regulate aging by acting as signaling molecules (e.g. as nuclear hormone receptor ligands). Indeed, the lipid profiles in a number of species change as animals age resulting in the acceleration of cell and tissue damage such that levels of certain lipids can serve as aging biomarkers (Papsdorf and Brunet, 2019). Also, studies have reported that fatty acids can alter animal physiology through the modification of chromatin states either by the direct addition of lipid moieties to chromatin or by activating signaling pathways that affect chromatin landscapes (Hansen et al. 2013; Papsdorf and Brunet, 2019). Furthermore, MUFAs have been shown to be the link between H3K4me3 modifiers and lifespan in *C. elegans*, while the inheritance of epigenetic memory through soma-germline crosstalk in response to cellular stresses are associated with the H3K4me3 modification complex (Tauffenberger and Parker, 2014; Kishimoto et al. 2016; Han et al. 2017). Altogether, these studies provide an exciting groundwork to examine how the chromatin states in PD_Stv_ animals correlate with metabolic plasticity due to early life starvation across generations.

During the course of its natural history, *C. elegans* occupies ephemeral environments such as rotting fruit or decomposing vegetation, where conditions and food availability are highly unpredictable. The dauer larva affords *C. elegans* a survival and dispersal strategy to escape harsh environmental conditions by often associating with passing invertebrate carriers. Accordingly, *C. elegans* is often found in the wild as dauers. Once a food source is found, dauers resume reproductive development to colonize the new habitat. Upon exhaustion of resources and population expansion, young larvae enter dauer and thereby repeating the boom and bust life cycle (Schulenburg and Félix, 2017). Because of frequent environmental perturbations, an adopted phenotypic plasticity strategy would ensure an advantage in species survival. The generation following a bust period would inherit the cellular programs for increased somatic lipid stores. It is thus remarkable that the cellular mechanisms to ensure survival of the species are fundamentally similar between humans and nematodes, two species that have diverged hundreds of millions of years ago, once again underscoring the relevance of a simple roundworm in understanding basic animal physiology.

## Acknowledgments

We are grateful for the generous gift of dafachronic acid from Frank Schroeder and Pooja Gubibanda for advice on its use. We thank Leszek Kotula and Angelina Regua for access to their microscope, and Eleanor Maine for thoughtful comments on this manuscript. Strain BS1080 was kindly provided by Tim Schedl. This work was partially supported by a NIHR01GM129135 grant to S.E.H. We thank the CGC, which is funded by the NIH Office of Research Infrastructure Programs (P40 OD010440), for providing strains.

## Author contributions

M.C.O. and S.E.H. conceived and designed experiments. M.C.O. and A.M.N. performed experiments. M.C.O., A.M.N., and S.E.H. analyzed the data. M.C.O. and S.E.H. wrote the manuscript.

## STAR ★ METHODS

Detailed methods are provided in the online version of this paper and include the following:

- KEY RESOURCES TABLE
- CONTACT FOR REAGENT AND RESOURCE SHARING
- EXPERIMENTAL MODEL AND SUBJECT DETAILS

- *C. elegans* strains and husbandry
- METHOD DETAILS

- Plasmid construction and transgenic strains
- Brood assays
- Oil Red O staining
- RNA interference
- Dafachronic acid supplementation
- Oleic acid supplementation
- Germ cell rows
- RNA extraction
- Quantitative reverse transcription real-time PCR
- QUANTIFICATION AND STATISTICAL ANALYSIS
- DATA AND SOFTWARE AVAILABILITY

### CONTACT FOR REAGENT AND RESOURCE SHARING

Further information and requests for resources and reagents should be directed to and will be fulfilled by the Lead Contact, Sarah E. Hall.

### EXPERIMENTAL MODEL AND SUBJECT DETAILS

#### *C. elegans* strains and husbandry

N2 Bristol wild-type strain was used as the reference strain. Worms were grown at 20°C unless otherwise indicated in Nematode Growth Medium (NGM) seeded with *Escherichia coli* OP50 using standard methods (Brenner et al. 1974; Stiernagle 2006). Mutants that were not previously backcrossed were backcrossed at least four times to our laboratory N2 wild-type before use. All strains used in this study are listed in the Key Resources Table.

### METHOD DETAILS

#### Plasmid construction and transgenic strains

To clone P*glp-4::glp-4 cDNA::gfp::glp-4* 3’UTR, primers sets MO2454 & MO2455; MO2516 & MO2507; MO2508 & MO2468; MO2469 & MO2470; and MO2471 & MO2472 were used to amplify pUC19, 1.2 kb of the *glp-4* promoter, the *glp-4* cDNA (using a cDNA library prepared from total RNA of a mixed N2 population as template), the *gfp* gene (using Fire vector pPD95.75 as the template), and 500 bp of the *glp-4* 3’UTR, respectively, using Phusion DNA polymerase (NEB). Assembly of PCR products was done using the NEBuilder HiFi DNA Assembly (E2621) to generate pMO555. P*glp-4::glp-4 cDNA::gfp::glp-4* 3’UTR was cloned into the pCFJ151 MosSCI vector to generate plasmid pMO557 using primer sets MO2499 & MO2500 and MO2514 & MO2504 to amplify pCFJ151and P*glp-4::glp-4 cDNA::gfp::glp-4* 3’UTR (using pMO555 as template), respectively, using Phusion DNA polymerase (NEB) and assembled with the NEBuilder HiFi DNA Assembly. To clone P*nhx-2::glp-4 cDNA::gfp::glp-4* 3’UTR, overlap extension PCR was done with primers MO2544 (*Xma*I) and MO2533 (with genomic DNA as template); MO2534 and MO2532 (*Sbf*I) (with pMO555 as template). The resulting amplicon was digested with *Xma*I and *Sbf*I and cloned into pUC19 to generate pMO562. P*nhx-2::glp-4 cDNA::gfp::glp-4* 3’UTR was cloned into pCFJ151 using the NEBuilder HiFi DNA Assembly and PCR products from primers MO2509 & MO2504 (with pMO562 as template) and primers MO2499 & MO2500 (with pCFJ151 as template) to create pMO565. To clone P*pgl-1::glp-4 cDNA::gfp::glp-4* 3’UTR, overlap extension PCR was done with primers MO2545 (*Xma*I) and MO2536 (with genomic DNA as template); MO2537 and MO2532 (*Sbf*I) (with pMO555 as template). The resulting amplicon was digested with *Xma*I and *Sbf*I and cloned into pUC19 to generate pMO559. P*pgl-1::glp-4 cDNA::gfp::glp-4* 3’UTR was cloned into pCFJ151 using the NEBuilder HiFi DNA Assembly and PCR products from primers MO2510 & MO2504 (with pMO559 as template) and primers MO2499 & MO2500 (with pCFJ151 as template) to create pMO566. To clone P*zfp-2::glp-4 cDNA::gfp::glp-4* 3’UTR, overlap extension PCR was done with primers MO2546 (*Xma*I) and MO2538 (with genomic DNA as template); MO2539 and MO2532 (*Sbf*I) (with pMO555 as template). The resulting amplicon was digested with *Xma*I and *Sbf*I and cloned into pUC19 to generate pMO560. P*zfp-2::glp-4 cDNA::gfp::glp-4* 3’UTR was cloned into pCFJ151 using the NEBuilder HiFi DNA Assembly and PCR products from primers MO2511 & MO2504 (with pMO560 as template) and primers MO2499 & MO2500 (with pCFJ151 as template) to create pMO567. Plasmids pMO557, pMO565, pMO566, and pMO567 were used to generate single copy insertions, *pdrSi1* (*Pglp-4::glp-4 cDNA::gfp::glp-4 3’UTR*), *pdrSi2* (*Pnhx-2::glp-4 cDNA::gfp::glp-4 3’UTR*), *pdrSi3* (*Pzfp-2 ::glp-4 cDNA::gfp::glp-4 3’UTR*), and *pdrSi4* (*Ppgl-1::glp-4 cDNA::gfp::glp-4 3’UTR*) by Mos1-mediated single copy insertion (MosSCI) (Frokjaer-Jensen et al. 2008). All primers sequences are listed in the Key Resources Table. Following integration into *unc-119(ed3)* animals segregated from EG6699 [*ttTi5605 II; unc-119(ed3) III; oxEx1578* (*eft-3p::gfp + Cbr-unc-119*)] that have lost the *oxEx1578* array, MosSCI insertions were genetically crossed into the *glp-4(bn2)* background.

#### Brood assays

Ten L4 larvae were placed individually onto 35mm NGM plates seeded with *E. coli* OP50 and incubated at 20°C. Animals were transferred daily to fresh 35mm NGM plates until egg laying ceased. Live progeny from each egg laying plate were counted. Assays were done from at least three biological independent replicates.

#### Oil Red O staining

Fat stores were stained using Oil Red O (ORO) dye as described by O’Rourke et al. 2009. Age matched one-day-old adults were washed from 60mm seeded NGM plates with 1x PBS pH 7.4 and rinsed 3-4 times until they were cleared of bacteria. Worms were permeabilized in 1x PBS pH 7.4 with an equal volume of 2x MRWB buffer (160 mM KCl, 40 mM NaCl, 14 mM Na_2_EGTA, 1 mM spermidine-HCl, 0.4 mM spermine, 30 mM PIPES pH 7.4, 0.2% β-mercaptoethanol) and supplemented with 2% paraformaldehyde. Samples were rocked for 1 hour at room temperature. Following fixation, worm samples were washed with 1x PBS pH7.4, resuspended in 60% isopropanol, and incubated at room temperature for 15 minutes. An Oil Red O (ORO) stock solution (prepared beforehand as a 0.5 g/100 mL in isopropanol and equilibrated in the dark at room temperature at least several days) was diluted to 60% with dH_2_O and rocked for at least one day to be used as the working stock. The ORO working stock was filtered through a 0.22 um filter immediately before use. Fixed worms were incubated in filtered ORO working stock and rocked overnight at room temperature. Next day, worm samples were allowed to settle and the ORO stain was removed. Worm pellets were washed once with 1x PBS pH7.4 and resuspended in 200 μL of 1x PBS with 0.01% Triton X-100. Aliquots of worm samples were mounted onto microscope glass slides and imaged.

Images were captured with either a Nikon Eclipse Ci with Spot 5.2 software or with an iPhone through iDu Optics*^®^* equipped with a LabCam*^®^* adapter (New York, NY, USA). Color images were separated into their RGB channel components and the intensity measured on the green channel (Yen et al. 2010) using ImageJ (NIH).

#### RNA interference

Gravid adults were treated with hypochlorite to obtain embryos using standard methods (Stiernagle 2006). Embryos were placed on NGM plates supplemented with 1mM IPTG and 50 μg/ml carbenicillin seeded with a 10x concentrated bacterial culture expressing the *k09f5.2* (*vit-1*) RNAi clone obtained from the Ahringer library (Kamath et al. 2001). Embryos were allowed to grow until adulthood at which time they were treated again with hypochlorite to obtain embryos. The recovered embryos were grown until day one of adulthood and collected for ORO staining.

#### Dafachronic acid supplementation

The Δ^7^ form of dafachronic acid (Δ^7^-DA) (a kind gift from Frank Schroeder) was added to a freshly grown culture of *E. coli* OP50 at a concentration of 40 nM and immediately seeded onto NGM plates. NGM plates supplemented with an equivalent volume of ethanol (the Δ^7^-DA solvent) to those of the Δ^7^-DA-supplemented NGM plates were used as the control plates. Seeded plates were allowed to dry overnight before use.

#### Oleic acid supplementation

Animals were induced into dauer by starvation on peptone-less NGM plates seeded with *E. coli* OP50 pre-loaded with oleic acid (NuChek Prep, Inc.; Elysian, Minnesota, USA) as described by Devkota et al. 2017. OP50 was grown overnight at 37°C in liquid LB supplemented with 600 μM of oleic acid or with an equivalent volume of ethanol (the oleic acid solvent) to serve as the control. Cultures were pelleted and washed several times with M9 buffer (Stiernagle, 2006) and resuspended at a 10x concentration. The 10x OP50 was seeded onto peptone-less NGM plates and allowed to dry overnight before use.

#### Germ cell rows

All worm strains used for germ cell row counting have the integrated transgene *gld-1*::*gfp*::*3xFLAG* from the BS1080 strain in their genetic background. Worms with L3 vulva morphology were identified by using Nomarski DIC microscopy at 630x (Seydoux et al. 1993) and DAPI stained using the standard whole worm DAPI staining protocol. The stained worms were imaged using a Leica DM5500B microscope with a Hamamatsu camera controller C10600 ORCA-R2. When performing the germ cell row counts, the start of the transition was identified when at least two cells in a row exhibited the crescent-shaped nuclei morphology (Shakes et al. 2009).

#### RNA extraction

Total RNA was extracted using TRIzol Reagent (Life Technologies). Four volumes of TRIzol reagent were added to a frozen worm pellet followed by vigorous vortexing for 20 minutes. Samples were centrifuged in a tabletop centrifuge at maximum speed and the cleared supernatant was transferred to a fresh tube. The RNA was precipitated with equal volume of isopropanol at −80°C for at least 30 minutes. RNA pellets were washed with cold 75% ethanol, dried, and resuspended in RNase-free water.

#### Quantitative reverse transcription real-time PCR

Total RNA was treated with DNaseI (NEB) and processed with Superscript IV First Strand Synthesis Systems (Life Technologies) using oligo (dT) primers following the recommendations of the manufacturer. Real-time PCR was done with iTaq Universal SYBR Green Supermix (BioRad) according to the recommendations of the manufacturer. C_t_ normalization was done using *act-1*. All primer sequences are listed in Key Resources Table.

### QUANTIFICATION AND STATISTICAL ANALYSIS

Statistical significance was determined using Prism 8 (GraphPad Software) using the statistical tests indicated in the figure legends.

### DATA AND SOFTWARE AVAILABILITY

Not applicable.

## Supplemental figure legends

**Figure S1.**
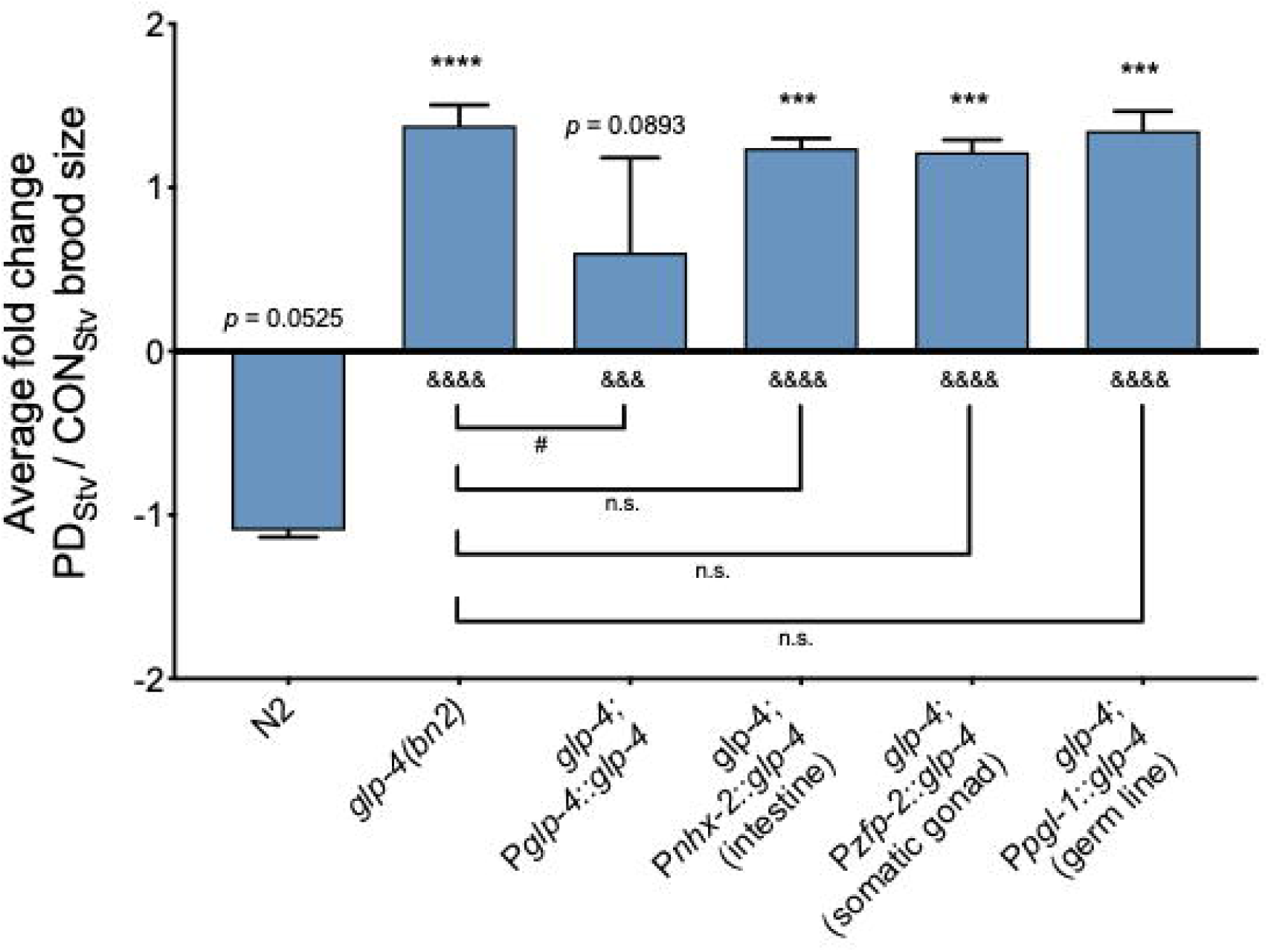
GLP-4 is required in multiple tissues to regulate adult reproductive plasticity. Brood size comparison of PD_Stv_ relative to CON_Stv_ in wild-type N2, *glp-4(bn2)*, and tissue-specific rescue strains of *glp-4*. Assays were done at 15°C from at least three biological independent trials. *** *p* < 0.001, **** *p* < 0.0001, and specified *p*-value (Student’s *t*-test) refer to the comparison of PD_Stv_ to CON_Stv_ of a given genotype. ^&&&^ *p* < 0.001 and ^&&&&^ *p* < 0.0001 (one-way ANOVA with Fisher’s LSD test) refer to the PD_Stv_/CON_Stv_ brood size change between a given mutant genotype to wild-type N2. ^#^ *p* < 0.05 and n.s. (one-way ANOVA with Fisher’s LSD test) refer to the PD_Stv_/CON_Stv_ brood size change between *glp-4(bn2)* and the indicated *glp-4* tissue-specific MosSCI rescues. Additional brood size data is provided in Supplemental Table S11.

**Figure S2.**
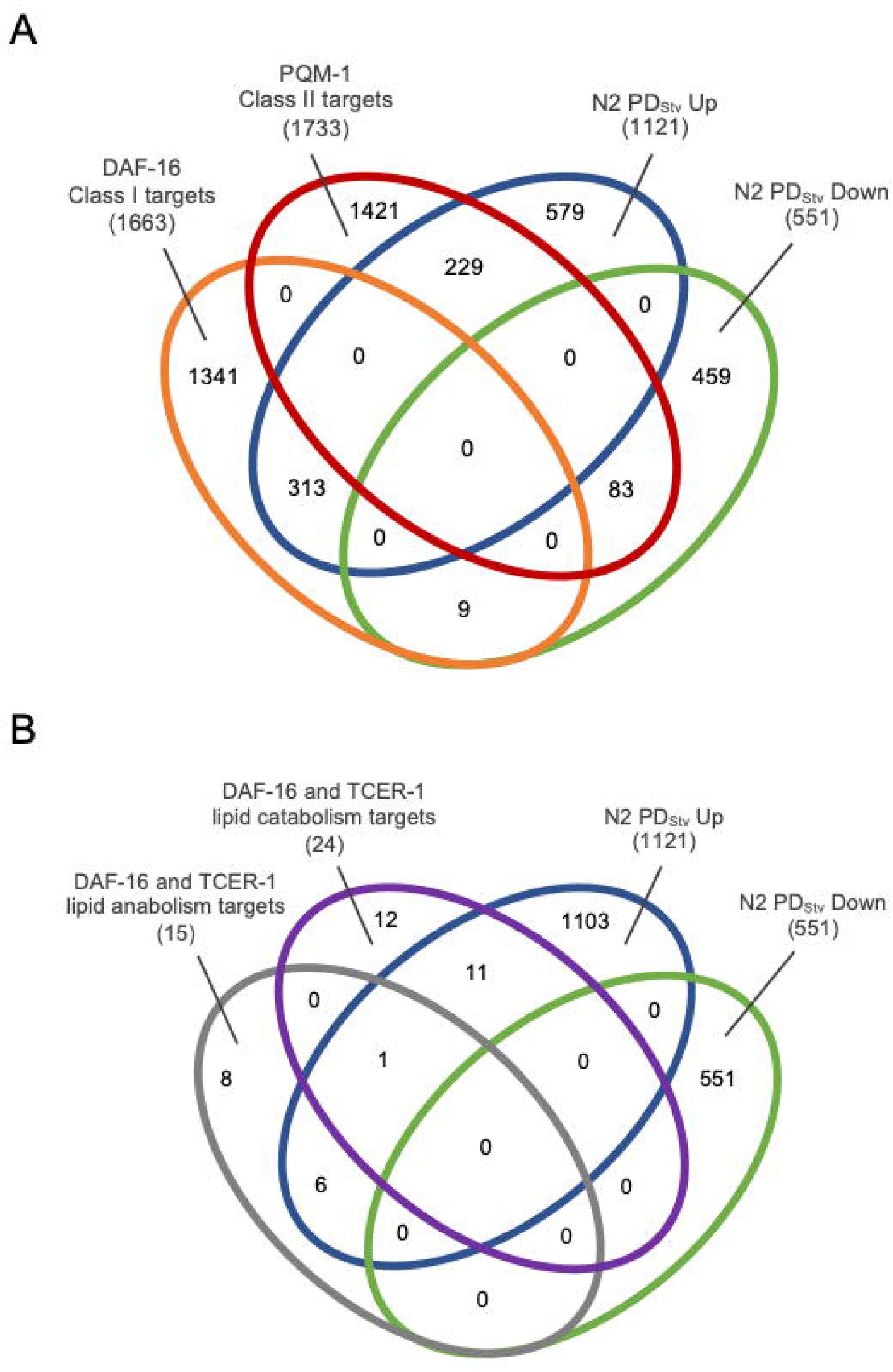
Differentially expressed N2 PD_Stv_ genes are enriched for DAF-16 targets. (*A*) Venn diagram depicting overlap of DAF-16 and PQM-1 targets (Tepper et al. 2013) with differentially expressed N2 PD_Stv_ genes compared to CON_Stv_ adults (Ow et al. 2018). See Supplemental Table S3. (*B*) Venn diagram comparing DAF-16 and TCER-1 targets with functions in lipid metabolism from germ line-less *glp-1(e2141ts)* animals (Amrit et al. 2016) with differentially expressed N2 PD_Stv_ genes compared to CON_Stv_ adults (Ow et al. 2018). See Supplemental Table S5.

**Figure S3.**
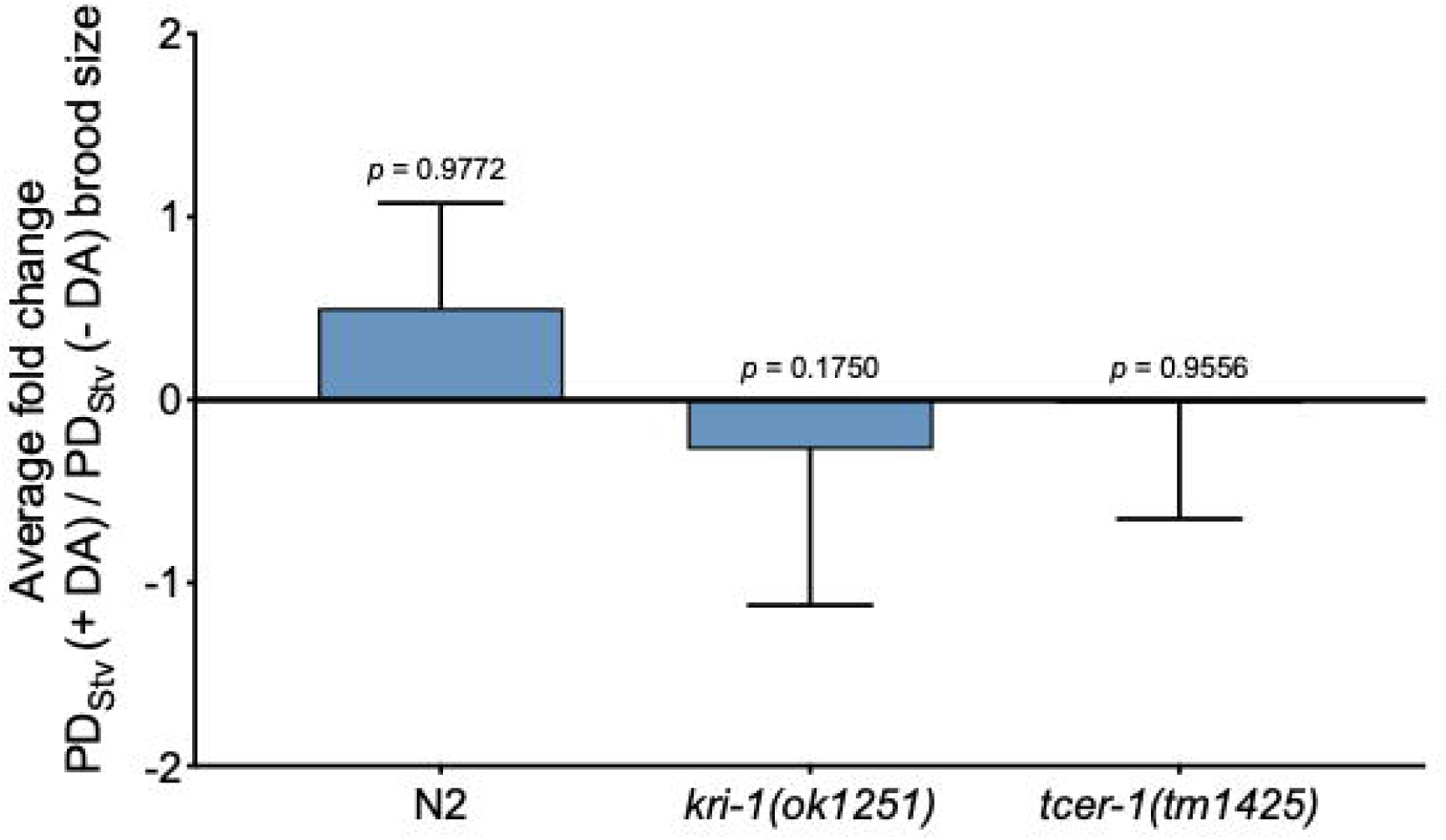
Brood size comparison of PD_Stv_ grown in the presence of 40 nM of Δ^7^-dafachronic acid (Δ^7^-DA) relative to the brood size of PD_Stv_ grown in the absence of Δ^7^-DA of wild-type N2, *kri-1(ok1251)*, and *tcer-1(tm1425)* strains. *p*-values (Student’s *t*-test) indicate the comparison between PD_Stv_ with Δ^7^-DA to PD_Stv_ without Δ^7^-DA of a given genotype. Assays were done at 20°C from three independent biological replicates. Additional brood size data is provided in Supplemental Table S12.

**Figure S4.**
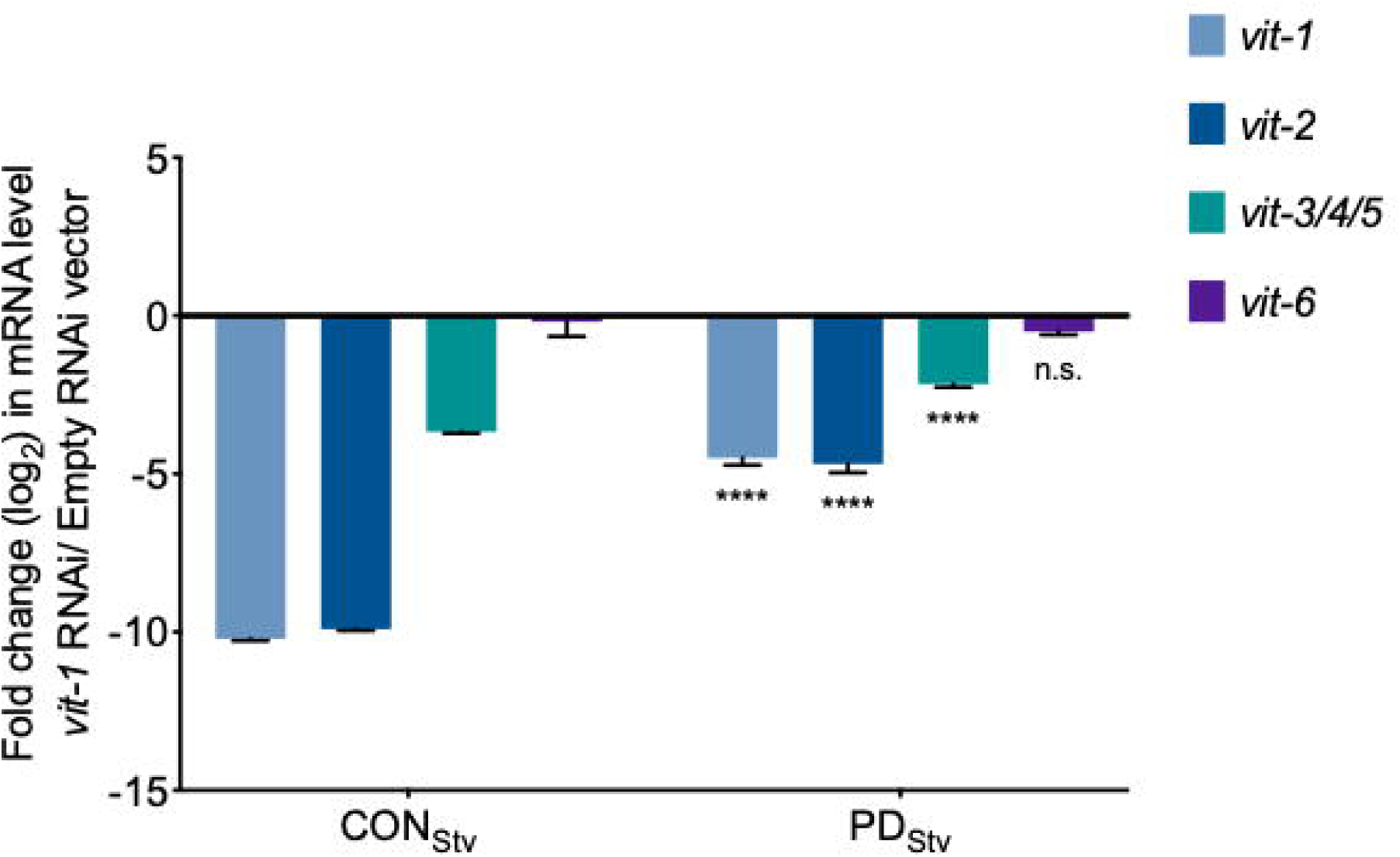
Reverse transcription quantitative PCR (qRT-PCR) measurements of the of the levels of *vit-1*, *vit-2*, *vit-3/4/5*, and *vit-6* mRNA in one-day-old-adults following RNAi-by-feeding of *k09f5.2* (*vit-1*). *act-1* was used as the normalization control. **** *p* < 0.0001 and n.s. (no significance) (Student’s *t*-test) refer to the comparison of PD_Stv_ to CON_Stv_ for a given RNAi knockdown. Primer sequences are provided in Supplemental Table S16.

**Figure S5.**
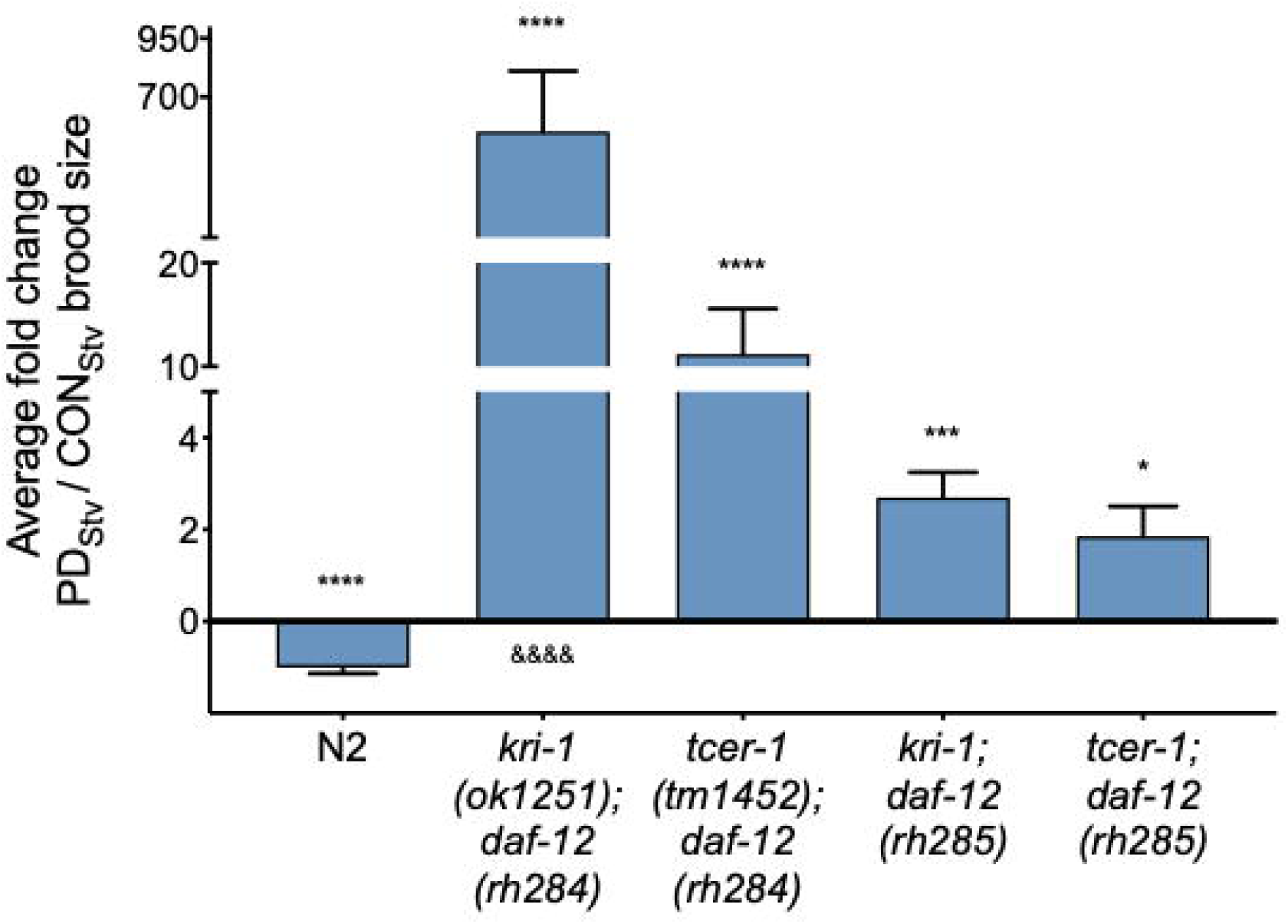
DAF-12 acts downstream of TCER-1 and KRI-1. Brood size comparison of PD_Stv_ relative to CON_Stv_ for wild-type N2, *kri-1(ok1251); daf-12(rh284)*, *tcer-1(tm1425); dat-12(rh284)*, *kri-1(ok1251); daf-12(rh285)*, and *tcer-1(tm1425); dat-12(rh285)*. * *p* < 0.05; *** *p* < 0.001, **** *p* < 0.0001 (Student’s *t*-test) indicates the brood size comparison between PD_Stv_ relative to CON_Stv_ of a given genotype. ^&&&&^ *p* < 0.0001 (one-way ANOVA with Fisher’s LSD test) indicates the comparison of PD_Stv_/CON_Stv_ brood size change between a given genotype to wild-type N2. Assays were performed at 20°C from at least three independent biological replicates. Additional brood size data is provided in Supplemental Table S10.

**Figure S6.**
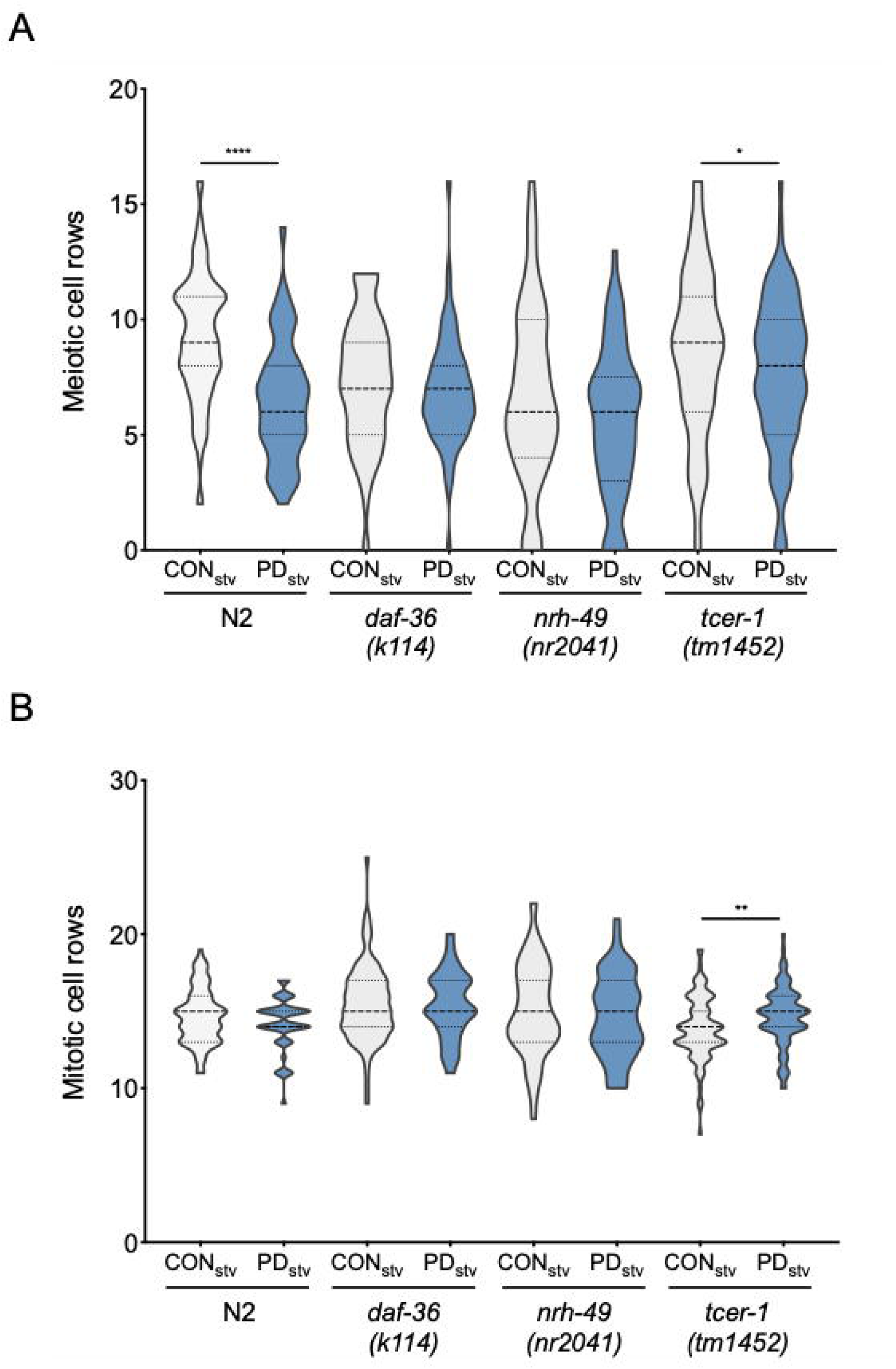
DAF-36 and TCER-1 are required for the delay in the onset of proliferation in animals that have experienced starvation-induced dauer formation. (*A*) The number of meiotic (transition zone) cell rows in CON_Stv_ and PD_Stv_ wild-type N2, *daf-36(k114), nhr-49(nr2041)*, and *tcer-1(tm1452)* at L3 stage was determined by counting the germ cells rows that express the crescent shaped morphology in DAPI stained worms. (*B*) The number of mitotic cell rows in CON_Stv_ and PD_Stv_ wild-type N2, *daf-36(k114), nhr-49(nr2041)*, and *tcer-1(tm1452)* at L3 stage was determined by subtracting the meiotic cell rows from the total cell rows for each animal. Student’s *t*-test comparison between CON_Stv_ and PD_Stv_ of the same genotype: * *p* <0.05, ** *p* < 0.01, **** *p* < 0.0001. Sample size is > 51 from at least three independent trials.

**Figure S7.**
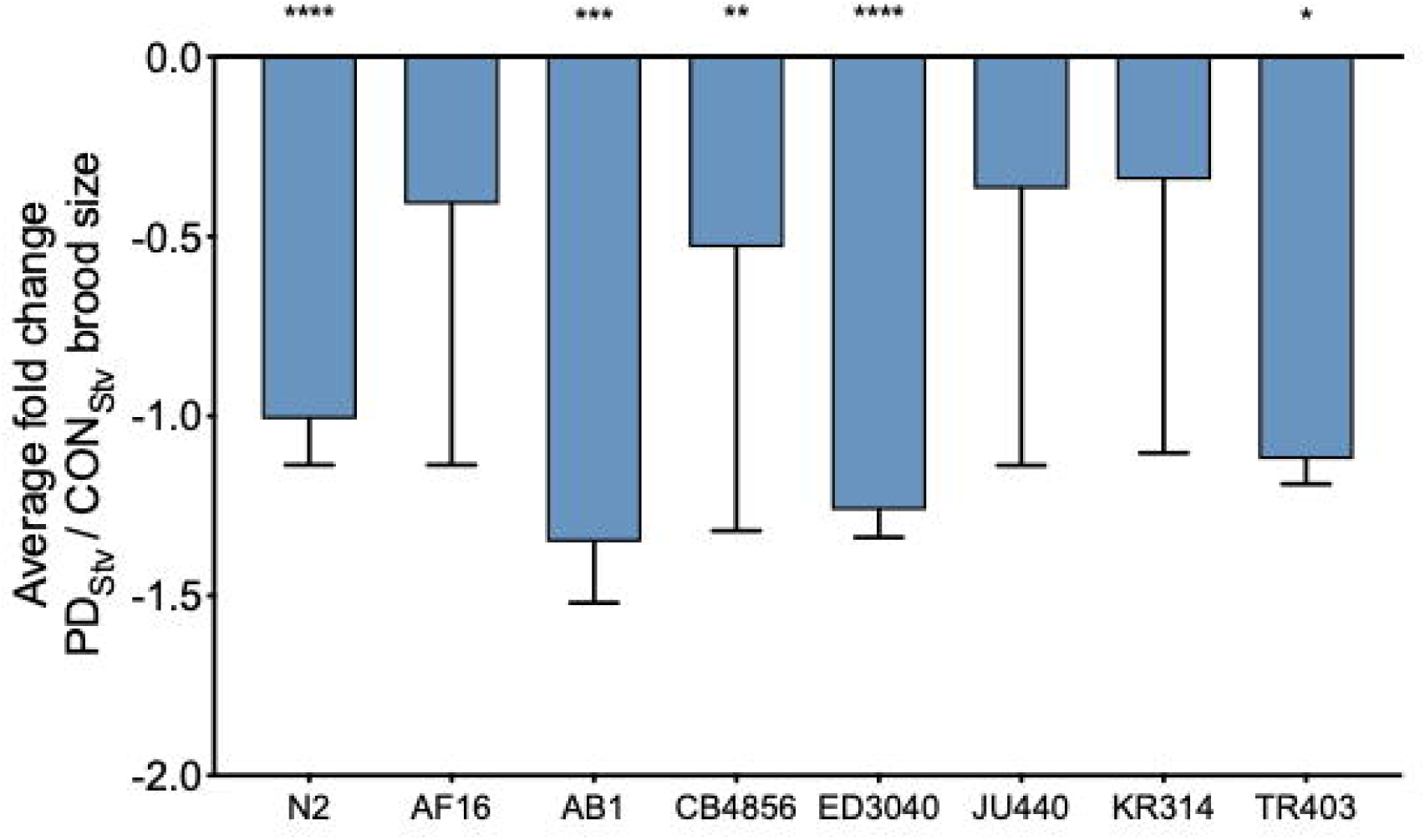
Wild *C. elegans* isolates exhibit adult reproductive plasticity. Brood size comparison of PD_Stv_ relative to CON_Stv_ for *C. elegans* N2 Bristol, *C. briggsae* AF16, *C. elegans* AB1, CB4856, ED3040, JU440, KR314, and TR403. * *p* < 0.05; ** *p* < 0.01, *** *p* < 0.001, **** *p* < 0.0001 (Student’s *t*-test) indicate the brood size comparison between PD_Stv_ relative to CON_Stv_ of a given genotype. Assays were performed at 20°C from at least three biologically independent replicates. Additional brood size data is provided in Supplemental Table S13.

