## Supplemental Material for "Somatic aging pathways regulate reproductive plasticity in *Caenorhabditis elegans*"

### Supplemental Results and Discussion

*GLP-4 is required globally to affect reproductive plasticity resulting from early life starvation*

We have previously shown that postdauer adults exhibit gene expression and reproductive phenotypes that reflect their environmental and developmental history. In adults that experienced starvation-induced dauer formation ($PD_{Stv}$), we showed that gene expression changes of somatically-expressed seesaw genes and their characteristic reduced brood size relative to control adults ($CON_{Stv}$) required a functional *glp-4* gene (Ow et al. 2018). The *glp-4* gene encodes a valyl aminoacyl tRNA synthetase that is expressed in the intestine, somatic gonad, and the germ line (Rastogi et al. 2015). The temperature-sensitive *bn2* allele is a partial loss-of-function lesion that likely results in decreased translation in both somatic and germline tissues (Beanan and Strome 1992; Rastogi et al. 2015). *glp-4(bn2)* mutants exhibit increased adult lifespan and stress resistance that require crosstalk between the germ line and the soma via endocrine signaling pathways (Arantes-Oliveira et al. 2002).

To address whether the expression of *glp-4* in the intestine, somatic gonad, or the germ line affect reproductive plasticity between control and $PD_{Stv}$ adults, we constructed single copy rescuing transgenes with tissue specific promoters using Mos1-mediated single copy insertion (MosSCI). We performed brood size assays of control and $PD_{Stv}$ adults at 15ºC, the permissive temperature at which *glp-4(bn2)* is fertile. While the brood size of $PD_{Stv}$ wild-type N2 adults was decreased to near significance (*p* = 0.0525) at the low temperature, the brood size of *glp-4(bn2)* $PD_{Stv}$ adults increased significantly, consistent with our previous observation (Supplemental Fig. S1; Supplemental Table S11) (Ow et al. 2018). Expression of *glp-4* under its endogenous promoter partially rescued the increased brood size of $PD_{Stv}$ *glp-4* adults (Supplemental Fig. S1; Supplemental Table S11). However, limited expression of *glp-4* in the intestine, somatic gonad, or the germ line resulted in a brood size phenotype similar to that of *glp-4(bn2)* mutants (Supplemental Fig. S1; Supplemental Table S11). These results suggest that the contribution of GLP-4 to the reproductive plasticity of $PD_{Stv}$ adults is not due to its function in a singular tissue (germ line, somatic gonad or intestine) but rather from multiple locations to promote inter-tissue crosstalk.

*The set of genes with altered expression in PDstv adults is significantly enriched for DAF-16 Class I and Class II targets*

We observed that 18.82% (313 genes; $p < 2.52e\text{-}91$, hypergeometric test) of the reported 1,663 DAF-16 Class I target genes and 13.21% (229 genes; $p < 1.96e\text{-}37$) of the 1,733 PQM-1 Class II target genes significantly overlapped with the 1,121 somatically-enriched genes that we previously reported to be up-regulated in $PD_{Stv}$ adults. We also observed significant overlaps between our previously reported 551 down-regulated genes in $PD_{Stv}$ adults with DAF-16 Class I genes (0.54%; 9 genes; $p < 1.03e\text{-}11$, hypergeometric test) and PQM-1 Class II genes (4.79%; 83 genes; $p < 2.83e\text{-}07$) (Tepper et al. 2013; Ow et al. 2018) (Fig. S2A; Supplemental Table S3).

A subset of DAF-16 targets are genes with functions in fat metabolism. DAF-16, along with TCER-1, alter the expression of lipid biosynthesis, storage, and hydrolysis genes to promote adult longevity in animals lacking a germ line (Armit et al. 2016). We also found a significant enrichment DAF-16 and TCER-1 targeted genes in our set of genes with altered mRNA levels in PDstv adults. Specifically, 47% (7 out of 15) of DAF-16 and TCER-1 target genes predicted to regulate lipid synthesis and storage are up-regulated in $PD_{Stv}$ adults ($p < 6.86e\text{-}06$, hypergeometric test). Similarly, 50% (12 out of 24) of lipid hydrolysis genes targeted by DAF-16 and TCER-1 are up-regulated in $PD_{Stv}$ adults ($p < 1.16e\text{-}09$) (Amrit et al. 2016; Ow et al. 2018) (Fig. S2B; Supplemental Table S5). Additionally, 27% (4 out of 15) of lipid catabolism genes ($p < 0.008$) and 12% (9 out of 74) of lipid anabolism genes ($p < 0.021$) that were not identified as targets of DAF-16 or TCER-1 were up-regulated in $PD_{Stv}$ adults (Amrit et al. 2016; Ow et al. 2018). Notably, none of the genes down-regulated in $PD_{Stv}$ adults were represented by the DAF-16 and TCER-1 lipid metabolic target genes (Amrit et al. 2016; Ow et al. 2018) (Fig. S2B). This observation was also true for lipid anabolism and catabolism genes that are not targets of DAF-16 or TCER-1.

*DAF-12 likely acts downstream of TCER-1 and KRI-1 to regulate reproductive plasticity*

To further examine the genetic interactions of DAF-12 and TCER-1, we performed epistasis analysis by measuring the brood sizes of control and $PD_{Stv}$ adults in *kri-1(ok1251); daf-12(rh284)* and *tcer-1(tm1425); daf-12(rh284)* double mutants, and in *kri-1(ok1251); daf-12(rh285)* and *tcer-1(tm1425); daf-12(rh285)* double mutants. For all four double mutants, we continued to observe a significantly increased brood size in $PD_{Stv}$

adults compared to controls, similar to the *daf-12(rh284)* and *daf-12(rh285)* single mutants (Fig. 1B, 2B, S5; Supplemental Table S1; Supplemental Table S10). The combination of mutations between *kri-1(ok1251)* and *tcer-1(tm1425)* with the *daf-12(rh284)* allele synergistically exacerbated the fertility phenotype of $PD_{Stv}$ adults to a more than 600-fold average compared to each of the single mutants, suggesting that these pathways act in parallel (Fig. 1B, 2B, S5; Supplemental Table S1; Supplemental Table S10). However, the brood sizes of the *kri-1(ok1251); daf-12(rh285*) and *tcer-1(tm1425); daf-12(rh285)* double mutants were statistically indistinguishable from the *daf-12(rh285)* single mutant, instead suggesting that DAF-12 acts downstream of TCER-1 and KRI-1 in the same pathway (Fig. 1B, 2B, S5, Supplemental Table S1; Supplemental Table S10). Since the *daf-12* and *tcer-1* mutant strains used in our experiment are not null alleles, we must interpret these epistasis results cautiously. However, we can make some conclusions with respect to the phenotypes of *daf-12(rh284)* and *daf-12(rh285)*, which have previously been characterized for developmental timing and dauer formation defects (Antebi et al. 1998; 2000). The *daf-12(rh284)* mutant (P746S lesion in helix 12 of the LBD) displays delayed gonadal development while the *daf-12(rh285)* mutant (Q707stop mutation in the LBD after helix 9) has penetrant heterochronic phenotypes that include delayed gonadal and extragonadal developmental events (Antebi et al. 1998; 2000). In addition, our brood assay results show that the *rh285* allele has a more severe phenotype than *rh284* in terms of reproduction (Fig. 1B; Supplemental Table S1). These phenotypes are perhaps due to the differences in the nature of LBD disruption that have not been characterized. Although the epistasis results could be interpreted as DAF-12 acting in parallel to KRI-1 and TCER-1, we favor a model in which DAF-12 acts downstream of KRI-1 and TCER-1 due to the greater severity of the *daf-12(rh285)* allele over the *daf-12(rh284)* allele.

C. elegans *wild isolates exhibit reproductive plasticity*

The standard wild-type N2 strain was first cultivated in the laboratory over five decades ago, resulting the accumulation random mutations over thousands of generations in laboratory conditions atypical to what are experienced by natural populations (Sterken et al. 2015). We wondered whether the reproductive plasticity observed in the canonical laboratory wild-type N2 strain is an adaptive trait acquired over time due to adaptation of laboratory conditions, or represents a conserved mechanistic response to starvation stress like that observed for the inheritance of adiposity memory from $PD_{Stv}$ parents. We assessed the control and $PD_{Stv}$ brood sizes of six natural isolates representing various branches of the *C. elegans* phylogenetic tree (Andersen et al. 2012). We found that 4 out of 6 natural isolates (AB1, CB4856, ED3040, and TR403) displayed a significant decrease in fertility in $PD_{Stv}$ adults relative to controls (Fig. S7; Supplemental Table S13). The wild isolates JU440 and KR314 had a similar number of progeny between control and $PD_{Stv}$ adults (Fig. S7; Supplemental Table S13). Interestingly, when we measured the brood size in control and $PD_{Stv}$ adults of a nematode species closely related to *C. elegans*, *C. briggsae* AF16, it displayed a modest but statistically insignificant decrease in $PD_{Stv}$ brood size compared to control adults (Fig. S7; Supplemental Table S13). Thus, the adult reproductive plasticity between animals undergoing continuous development and those that have experienced dauer diapause as a result of starvation is a naturally occurring developmental trait and not an adaptive trait stemming from decades of cultivation in a laboratory.

Is the reproductive plasticity between $CON_{Stv}$ and $PD_{Stv}$ a purely hermaphroditic response to starvation? The male frequency in the hermaphroditic N2 is a low ~0.1% (Chasnov and Chow, 2002). We find that one natural isolate, CB4856, which harbors a higher frequency of males in their population than the laboratory N2 strain due to an increase of successful copulation events (Wegewitz et al. 2008), also exhibits $CON_{Stv}$/$PD_{Stv}$ reproductive plasticity (Fig. S7; Supplemental Table S13), suggesting that fertility plasticity between $CON_{Stv}$ and $PD_{Stv}$ may not be due to reduced male frequency. Because *C. elegans* is androdioecious (hermaphrodites and males), we cannot unequivocally state that reproductive plasticity is not a purely hermaphroditic response to starvation. It is possible that in dioecious species (females and males), the cellular mechanisms deployed in response to starvation-induced stress early in development are distinct from those in androdioecious species.
